# Role of Very Long-Chain Fatty Acyl-Ceramides in Endocytic Uptake

**DOI:** 10.64898/2026.11.02.759677

**Authors:** Isabel María Fernández-Golbano, Laia Huguet, Daniel Álvarez, Ian Garí, Oriol Trillo, Sergio Ivan Cheng Zhu, Elena Rebollo, María Isabel Geli

**Affiliations:** Institute for Molecular Biology of Barcelona (IBMB, CSIC), C/ Baldiri Reixac 15, 08028 Barcelona Spain; Materials and Sustainability Group, Department of Engineering, Universidad Loyola Andalucía, Avda. de las Universidades s/n. Dos Hermanas, 41704 Seville, Spain; Instituto de Investigación en Energía, Tecnología y Sostenibilidad (LETS), Universidad Loyola Andalucía, Seville, Spain

## Abstract

Specialized endoplasmic reticulum (ER) subdomains termed ER sterol exit sites (ERSES), transiently contact endocytic sites to locally supply sterols, which in turn initiate actin polymerization and drive membrane invagination. Osh2, an oxysterol-binding protein (OSBP)–related protein (ORP), mediates the non-vesicular sterol transfer at ER–endocytic contact sites (ER–ECS). In search for additional ERSES components, we identified Fat1, a fatty acid (FA) transporter and acyl-CoA synthetase with specificity for very long-chain (VLC) FAs (C20–C26). Analysis of VLC-FA anabolic and catabolic pathways revealed a specific requirement for VLC fatty acyl-ceramides (VLCA-Cers) in the constriction of endocytic invaginations. Strikingly, the Cer synthase physically interacted with the endocytic machinery, pointing to a direct role of VLCFA-Cers in membrane remodelling. Consistent with this view, molecular dynamics and live-cell imaging of a Cer probe indicated that the ORPs Osh1 and Osh3 contribute to Cer export from at ER-ECS and and that their depletion recapitulates the endocytic defects observed in the absence of VLCFA-Cer. Our findings establish that local transport of VLCA-Cers is required for endocytic budding and reveal ERSES as hubs where Cer and sterol biosynthesis and transport converge.

## Introduction

Studies on clathrin- and actin-dependent endocytic uptake in yeast have led to the identification of more than sixty proteins directly involved in cargo recruitment, membrane deformation, and vesicle scission. Those include clathrin and the cargo adaptors, which are first recruited to the plasma membrane (PM); followed by the WASP-Myo and scission modules (Idrissi and Geli, 2014; Kaksonen et al., 2003; Kaksonen et al., 2005). The, WASP-Myo module comprises the N-WASP homolog Las17 and the myosin-I Myo5, and induces transient assembly of an actin cap branched by the Arp2/3 complex, required to drive membrane invagination (Galletta et al., 2008; Idrissi et al., 2012; Kukulski et al., 2012a; Kukulski et al., 2012b; Sun et al., 2006). The ATPase of the myosin-I, which sits at the PM / invagination junction (PMIJ), and the assembly of the N-BAR domain-containing proteins Syp1 and Bzz1 at the invagination neck, are then required to initiate constriction (Idrissi et al., 2012; Idrissi and Geli, 2014). Vesicle scission is finally assisted by the F-BAR proteins Rvs161 and Rvs167, the dynamin Vps1 and actin (Idrissi and Geli, 2014; Kishimoto et al., 2011; Smaczynska-de et al., 2012).

Live-cell imaging and genetic approaches using conditional mutants have unequivocally demonstrated the direct and local role of all these proteins in endocytosis. In contrast, although reverse genetics early on suggested that sterols and sphingolipids play a role in endocytic uptake (Munn et al., 1999; Zanolari et al., 2000), robust demonstration of their direct function has been difficult. This is partly due to their rapid dynamics, which hinder imaging and fine functional analysis.

Our group discovered that in yeast, the cortical endoplasmic reticulum (cER) transiently contacts endocytic sites to supply sterols required to timely initiate actin polymerization and to drive membrane invagination and vesicle scission (Encinar Del Dedo et al., 2021; Encinar Del Dedo et al., 2017). We found that the ER subdomains that interact with endocytic sites, termed ERSES (ER Sterol Exit Sites), are enriched in a subset of sterol biosynthetic enzymes (Encinar Del Dedo et al., 2021) and that the oxysterol-binding protein (OSBP)–related protein (ORP) Osh2 interacts both with the sterol biosynthetic and endocytic machineries and mediates sterol transfer at ER–endocytic contact sites (ER-ECS) (Encinar Del Dedo et al., 2021). We also found that the physical interaction of the OSBP-related lipid transfer domain (ORD) of Osh2 with the ergosterol biosynthetic machinery is essential to support the capacity of Osh2 to extract sterols from ERSES and, consequently, for its endocytic function (Encinar Del Dedo et al., 2021). Together with the nanometric localization of sterols at the PMIJ (Encinar Del Dedo et al., 2021), the data provided robust evidence for a direct and localized role of sterols in endocytic uptake.

In the present work, we searched for additional ERSES components. We identified Fat1, a fatty acid (FA) transporter and acyl-CoA synthetase that preferentially activates very long– chain (VLC) FAs (VLCFAs) with 20 to 26 carbons (C20–C26) (Choi and Martin, 1999). We demonstrated that VLC fatty acyl-CoAs (VLCA-CoAs) are required to support endocytic uptake though their contribution to the synthesis of VLC fatty acyl-ceramides (VLCA-Cers). Interestingly, we also found that the Cer synthase (CerS) complex, composed of the catalytic subunits Lac1 and Lag1 and the regulatory subunit Lip1 (Guillas et al., 2001; Schorling et al., 2001; Vallee and Riezman, 2005), interacts both with the ERSES components and the endocytic myosin-I Myo5, as well as with the BAR domain-containing proteins Syp1, Bzz1 and Rvs167; pointing to a direct and local role of VLCFA-Cers in endocytosis. Consistent with this view, molecular dynamics (MD) and live-cell imaging experiments using a newly developed Cer probe indicated that Osh1 and Osh3 contribute to VLCFA-Cer transport at ER-ECS and that depletion of these ORPs recapitulate the endocytic phenotypes installed upon elimination of VLCFA-Cers.

## Results

### VLCA-CoAs Are Required for Endocytic Uptake

To identify components of the ERSES that might inform on their molecular function, we searched the *Saccharomyces Genome Database* (SGD) for proteins that interact with two or more of the known ERSES components, which are the C-8 sterol isomerase Erg2; the δ(24)-sterol C-methyltransferase Erg6; the lanosterol synthase Erg7; and the 3-keto sterol reductase Erg27 (Fig. S1A) (Encinar Del Dedo et al., 2021). The search identified Fat1 as an interactor of Erg6, Erg7 and Erg27 (Tarassov et al., 2008), (Athenstaedt et al., 1999). Fat1 is a VLCA-CoA synthetase and FA transporter with preference for VLCA (C20-C26) (Fig. 1A) (Choi and Martin, 1999). The interactions of Fat1 with Erg7, Erg6 and Erg27 were confirmed by immunoprecipitation from cER and PM-enriched cellular fractions (Fernandez-Golbano et al., 2014), from strains expressing Green Fluorescent Protein (GFP)-tagged Erg6, Erg7 or Erg27, and Fat1-HA (Haemaglutinin) (Fig. 1B).

**Figure 1.**
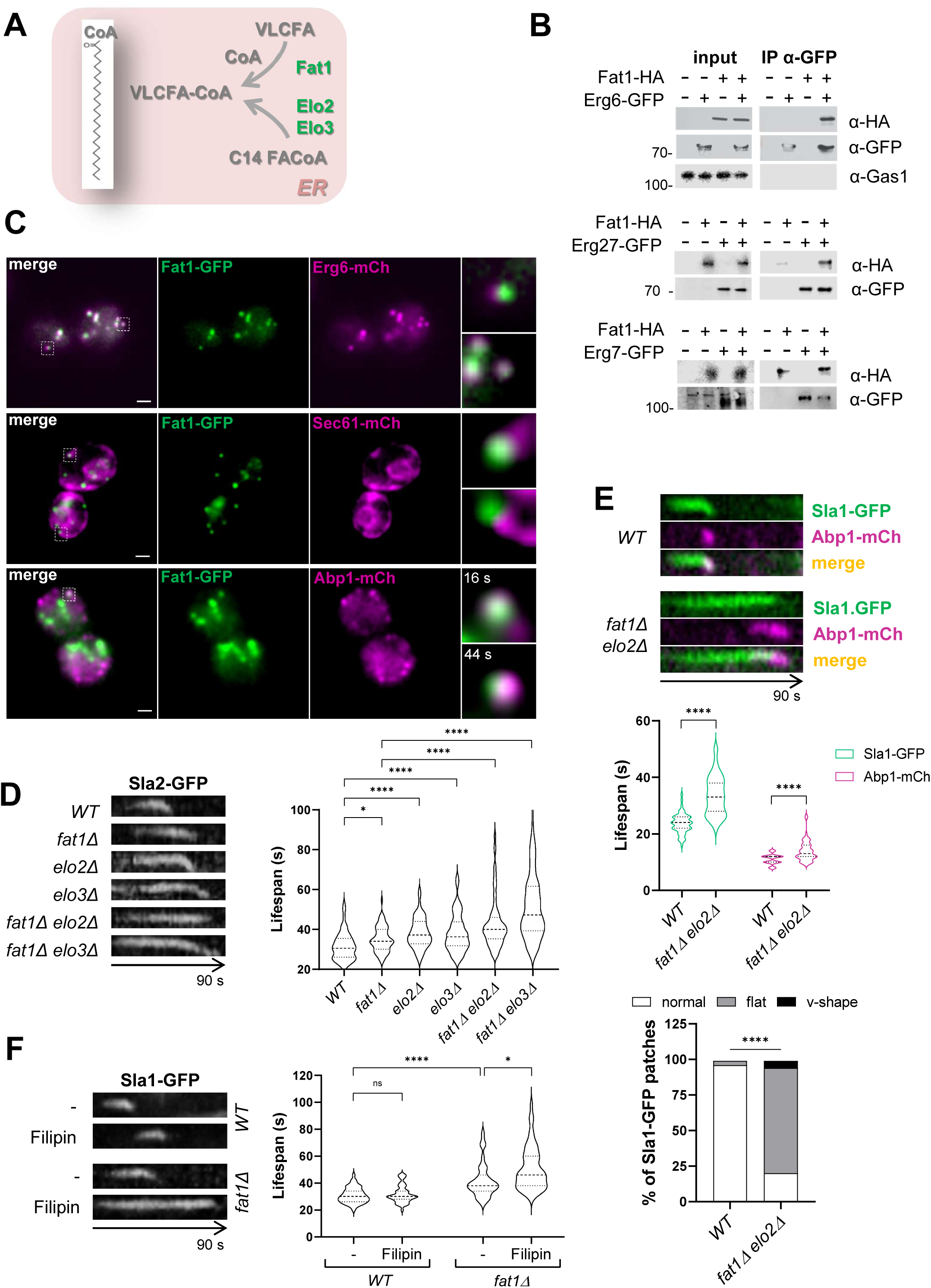
VLCFA-CoAs Are Required for Endocytic Uptake. **A.** Scheme depicting the pathways generating VLCFA-CoAs in yeast. **B.** Immunoblots decorated with anti-GFP (α-GFP) and anti-HA (α-HA) antibodies of anti-GFP agarose immunoprecipitates from non-denaturing protein extracts of yeast expressing the indicated Erg proteins tagged with GFP (+) or the untagged versions (-); and Fat1-HA (+) or the untagged protein (-). 10 µg of total protein from the extracts were loaded as input. **C.** Dual color epi-fluorescent micrographs of live yeast expressing Fat1-GFP and either Erg6-mCherry, Sec61-mCherry or Abp1-mCherry. Individual channels and the merge images are shown. Merged magnified micrographs of the depicted areas are shown on the right. Bar = 1 µm **D.** Kymographs of 2 s time-lapse epi-fluorescence movies of cortical Sla2-GFP patches from the indicated strains; and violin plots of the Sla2-GFP cortical patch lifespans. Asterisks indicate statistically significant differences between groups, determined by Kruskal-Wallis test with Dunn’s post-hoc correction for multiple comparisons. **E.** Kymographs of dual-color 2 s time-lapse fluorescence movies of cortical Sla1-GFP and Abp1-mCherry patches for the indicated strains; and violin plots of the cortical patch lifespans of each endocytic protein for the corresponding strains. Asterisks indicate statistically significant differences between groups determined by Mann-Whitney test. The bar plot indicated the percentage of patches showing abnormal flat or V-shaped kymographs representing endocytic coats that failed to internalize or undergo fission. Statistical significance was determined by chi-square test; asterisks indicate significant differences between groups. **F.** Kymographs of 2 s time-lapse epi-fluorescence movies of cortical Sla1-GFP patches for the indicated strains either treated with 5 µg/ml of filipin or mock treated; and violin plots of the Sla1-GFP cortical patch lifespan for the corresponding strains and experimental conditions. Asterisks indicate significant differences between groups determined by Kruskal-Wallis test with Dunn’s post-hoc correction for multiple comparisons. **D to F**. Dashed lines in the violin plots indicate the median and dotted lines, the interquartile range (25^th^–75^th^ percentiles). n > 50 endocytic patches from more than 20 cells for each strain and experimental condition, from a representative experiment.

To further investigate the possible localization of Fat1 at ERSES, we generated a strain expressing Fat1-GFP from the genomic locus and explored its co-localization with Erg6-mCherry. We have previously shown that Erg6 localizes at ERSES, as well as in Lipid droplets (LDs). LDs appear as large spherical organelles, which can be stained with the neutral lipid marker BODIPY and are normally associated with the nuclear envelop in our growth conditions (Encinar Del Dedo et al., 2021). ERSES instead are fainter cortical patches situated at the rims of the cER, which are hot spots for endocytosis (Encinar Del Dedo et al., 2021). As shown in figure 1C, Fat1-GFP and Erg6-mCherry fully co-localized or appeared closely apposed in large internal and fainter cortical structures. Co-expression of the ER marker Sec61-mCherry with Fat1-GFP demonstrated the presence of the faint cortical Fat1-GFP patches at the rims of the cER, where ERSES localize (Fig. 1C). Those cortical Fat1-GFP patches were hot spots for the assembly of endocytic actin, labeled with Abp1-mCherry (Fig. 1C, S1B and movie S1), as previously shown for ERSES (Encinar Del Dedo et al., 2021). These data confirmed that Fat1 is a *bona-fide* component of the ERSES.

Next, we investigated if Fat1 might be required for endocytic uptake by following the behaviour of the endocytic coat component Sla2-GFP, expressed from a centromeric plasmid. We have previously shown that in regions devoid of cER in a reticulon mutant, endocytic coats assemble but their cortical life span is longer and they fail to internalize (Encinar Del Dedo et al., 2017). Further, actin polymerization at endocytic sites is delayed and extended with respect to the coat recruitment (Encinar Del Dedo et al., 2017). A small but significant extension of the Sla2-GFP cortical patch life-span was observed in the *fat1Δ* strain, which was substantially exacerbated in combination with deletions of *ELO2* or *ELO3*, encoding elongases involved in *de novo* synthesis of VLCA-CoAs from C18 FA-CoA primers (Oh et al., 1997) (Fig. 1A, 1D). Analysis of endocytic uptake in the *fat1Δ elo2Δ* mutant using the membrane impermeable dye Lucifer yellow confirmed the endocytic defect in the cells lacking VLCA-CoAs (Fig. S1C). The *fat1Δ elo3Δ* strain exhibited slow growth and strong Lucifer yellow cytosolic staining (Fig. S1C), suggesting membrane permeability defects and therefore, we did not use it for further analyses.

To investigate in more detail the endocytic defect observed in the *fat1Δ elo2Δ* mutant, we generated a strain expressing the late coat Sla1-GFP marker from the endogenous *locus* and Abp1-mCherry, which decorates endocytic actin. Analysis of Sla1-GFP cortical dynamics relative to Abp1-mCherry confirmed an extended life-span of the endocytic coat at the PM and demonstrated an average 8 second (s) delayed onset of actin polymerization and an extended life-span of the endocytic actin marker Abp1 (Fig. 1E), similar to the phenotype observed in the *osh2Δ* mutant (Encinar Del Dedo et al., 2017). This suggested that VLCFA-CoAs might share an endocytic function with sterols. In accordance, treatment of the *fat1Δ* mutant with low concentrations of the sterol sequestering drug filipin-which do not alter endocytic uptake in a wild type (wt) strain-, exacerbated the endocytic phenotype of Fat1 depletion (Fig. 1F). However, in contrast to the *osh2Δ* mutant, where most Sla1-GFP patches finally move into the cytosol (Encinar Del Dedo et al., 2017), most of the Sla1-GFP cortical patches in the *fat1Δ elo2Δ* strain failed to internalize (Fig. 1E). This phenotype most closely recapitulated that observed in endocytic coats that do not get in touch with the cER in a reticulon mutant (Encinar Del Dedo et al., 2017), suggesting additional endocytic functions for VLCFA-CoAs, as compared to sterols.

### VLCA-Cers but not more Complex Sphingolipids are Required for Endocytic Uptake

Next, we asked which VLCA-CoAs anabolic or catabolic pathway(s) might be required for endocytic uptake. VLCA-CoAs are transported to peroxisomes for β-oxidation or they can be used to synthesize glycosylphosphatidylinositol (GPI)-anchors; Cers and more complex sphingolipids; a small fraction of phosphoinositides; or triacylglycerol and esterified sterols stored in LDs (Fig. 2A) (Erdbrugger and Frohlich, 2020). Depletion of the VLCA-CoA peroxisomal importer Pxa1 (van Roermund et al., 2012) or knock out of the gene encoding the β-oxidation entry enzyme Pox1 (Dmochowska et al., 1990), did not result in endocytic defects, as assessed by monitoring the Sla2-GFP cortical patch dynamics (Fig. 2B). To explore which anabolic pathway might then be required, we investigated the endocytic phenotype of the corresponding acyl-transferase mutants (Fig. 1A), including knock out strains for the genes encoding Gup1, involved in GPI-anchor synthesis (Bosson et al., 2006); Cst26 and its homolog Mlg1, which produce a small percentage of VLCA-phosphatidylinositols (Vazquez et al., 2016); a cuadruple knock out of the LD’s Acyl-CoA:sterol acyltransferases Are1 and Are2 and the triacylglycerol acyl-transferases Lro1 and Dga1, lacking LDs (Sandager et al., 2002); and a double mutant knock out of the genes encoding the essential Cer synthase (CerS) catalytic subunits Lac1 and Lag1, complemented with a cotton CerS catalytic subunit (GhLag1) that generates Cers with shorter acyl chains (C18) (Fig. 2A) (Epstein et al., 2012). Interestingly, we found that only the CerS mutant recapitulated the *fat1Δ elo2Δ* endocytic phenotype exhibiting both extended Sla2-GFP cortical patch lifespans (Fig, 2C), and inability to take up Lucifer Yellow (Fig. S2A). It is worth noticing that the *lac1Δ lag1Δ GhLAG1* strain still generates Cers and sphingolipids, but their acyl chains are shorter (Epstein et al., 2012), demonstrating the relevance of this particular lipid feature. Depletion of Isc1, a protein that generates Cers by hydrolysis of more complex sphingolipids (Sawai et al., 2000) did not cause an endocytic defect (Fig. S2B), indicating that endocytosis requires *de novo* synthesis of Cers rather than their catabolic production.

**Figure 2.**
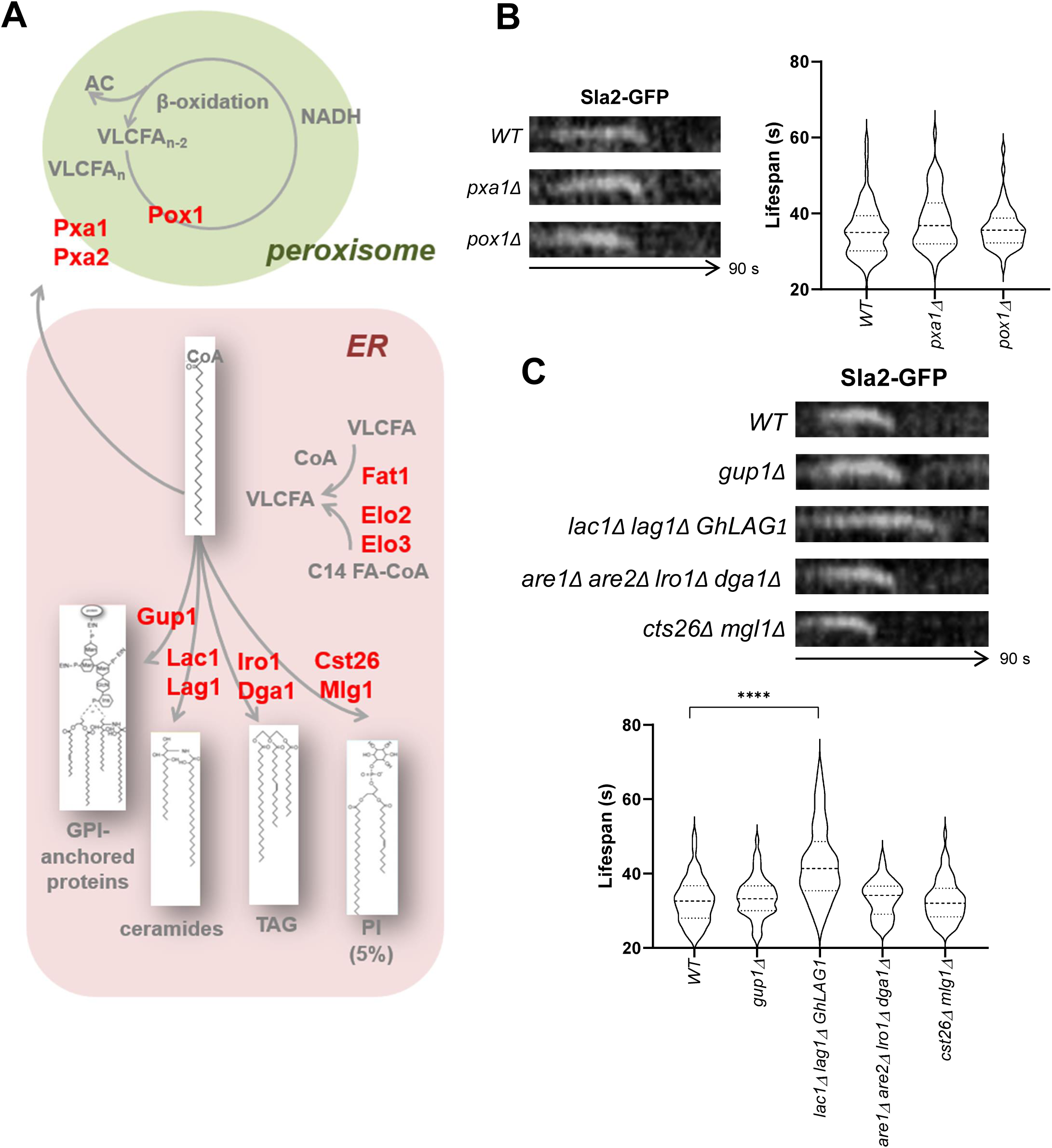
Synthesis of VLCA-Cers but not other VLCA-CoAs Anabolic or Catabolic Pathways Are Required for Endocytic Uptake. **A.** Scheme depicting the catabolic peroxisomal β-oxidation and the ER anabolic pathways for VLCA-CoAs in yeast. In red are depicted the enzymes or transporters involved. **B & C.** Kymographs of 2 ss time-lapse epi-fluorescence movies of cortical Sla2-GFP patches for the indicated strains; and violin plots of the Sla2-GFP cortical patch lifespans for the corresponding strains. Asterisks indicate statistically significant differences determined by Kruskal-Wallis test with Dunn’s post-hoc correction for multiple comparisons. Dashed lines in violin plots indicate the median and dotted lines the interquartile range (25^th^–75^th^ percentiles). n > 50 endocytic patches from more than 20 cells per strain, from a representative experiment.

Depletion of the human CerS2, which also generates VLCA-Cers, has been reported to install clathrin-mediated endocytic defects, indirectly through the miss-function of mitochondria, which accumulate ROS and secondarily down-regulates Hsc70, a chaperone involved in the disassembly of the clathrin coat (Volpert et al., 2017). Staining of wild type (wt) and *lac1Δ lag1Δ GhLAG1* yeast with DHE (dihydroethidium) (Liao et al., 2020) also revealed ROS accumulation in the absence of VLCFA-Cers in yeast (Fig. S2C). However, immunoblot analysis of the Ssa1-HA levels (the yeast Hsc70 homolog) did not indicate significant differences between the wt and the *lac1Δ lag1Δ GhLAG1* mutant (Fig. S2D). Further, depletion of the mitochondrial superoxide dismutase Sod2, which caused similar ROS levels, as compared with the CerS mutant (Fig. S2C), did not result in an endocytic phenotype (Fig. S2E), suggesting that high levels of ROS *per se* do impair endocytosis in yeast.

To confirm the requirement for Cer synthesis in endocytosis and explore the need for more complex sphingolipids, which are synthesized from Cers as they are transported to the Golgi, we assayed wt yeast treated with the CerS inhibitor Fumonisin B (Wu et al., 1995) and with Aureobasidin B, which inhibits Aur1, the phosphatidylinositol:Cer phosphoinositol transferase, which catalyzes the synthesis of inositol-phoshoCer (IPC) in the Golgi (Heidler and Radding, 1995) (Fig. 3A). As a positive control, we also assayed Myriocin, which targets the serine palmitoyltransferase (SPT) complex (Miyake et al., 1995) composed of Lcb1, Lcb2 and Tsc3; responsible for the condensation of serine with palmitoyl-CoA to form the 3-ketosphinganine long-chain base (Gable et al., 2000; Nagiec et al., 1994) (Fig 3A). Myriocin blocks endocytosis via the inhibition of the Pkh1/2 sphingoid based-activated pathway (Friant et al., 2001). As shown in figure 3B, treatment of cells with Fumonisin B recapitulated the *lac1Δ lag1Δ ghLAC1* mutant phenotype, further confirming the need for Cer synthesis in the process. The endocytic defect in cells treated with Myriocin was stronger than that of Fumonisin B, maybe reflecting a dual endocytic role of the SPT complex via the Pkh1/2-mediated sphingoid base signaling (Friant et al., 2001) and Cer synthesis. Interestingly though, treatment of cells with Aureobasidin A under conditions that prevent IPC synthesis (Reggiori and Conzelmann, 1998) did not alter endocytosis (Fig. 3B), suggesting that complex sphingolipids might not be required. To further test this hypothesis, we used a Tet-off Aur1 strain. Treatment of this strain under conditions reported to inhibit IPC synthesis (Yamaguchi et al., 2018), did not alter the dynamics of Sla2-GFP (Fig. 3C). In contrast, treatment of a Tet-off Lcb1 strain under the same conditions (Yamaguchi et al., 2018) significantly extended the Sla2-GFP cortical life-span, as seen for the Myriocin treated wt cells (Fig. 3B & C). Further confirming that complex sphingolipids are not required for endocytic uptake, we did not observe altered Sla2-GFP cortical patch dynamics in cells depleted of either the Mannosylinositol-phosphoryl-Cer (MIPC) synthase catalytic subunits Sur1 and Csh1, or the Inositol-phosphotransferase Ipt1, involved in synthesis of mannose-(inositol-P)_2_-Cer (M(IP)_2_C) (Fig. 3A & D).

**Figure 3.**
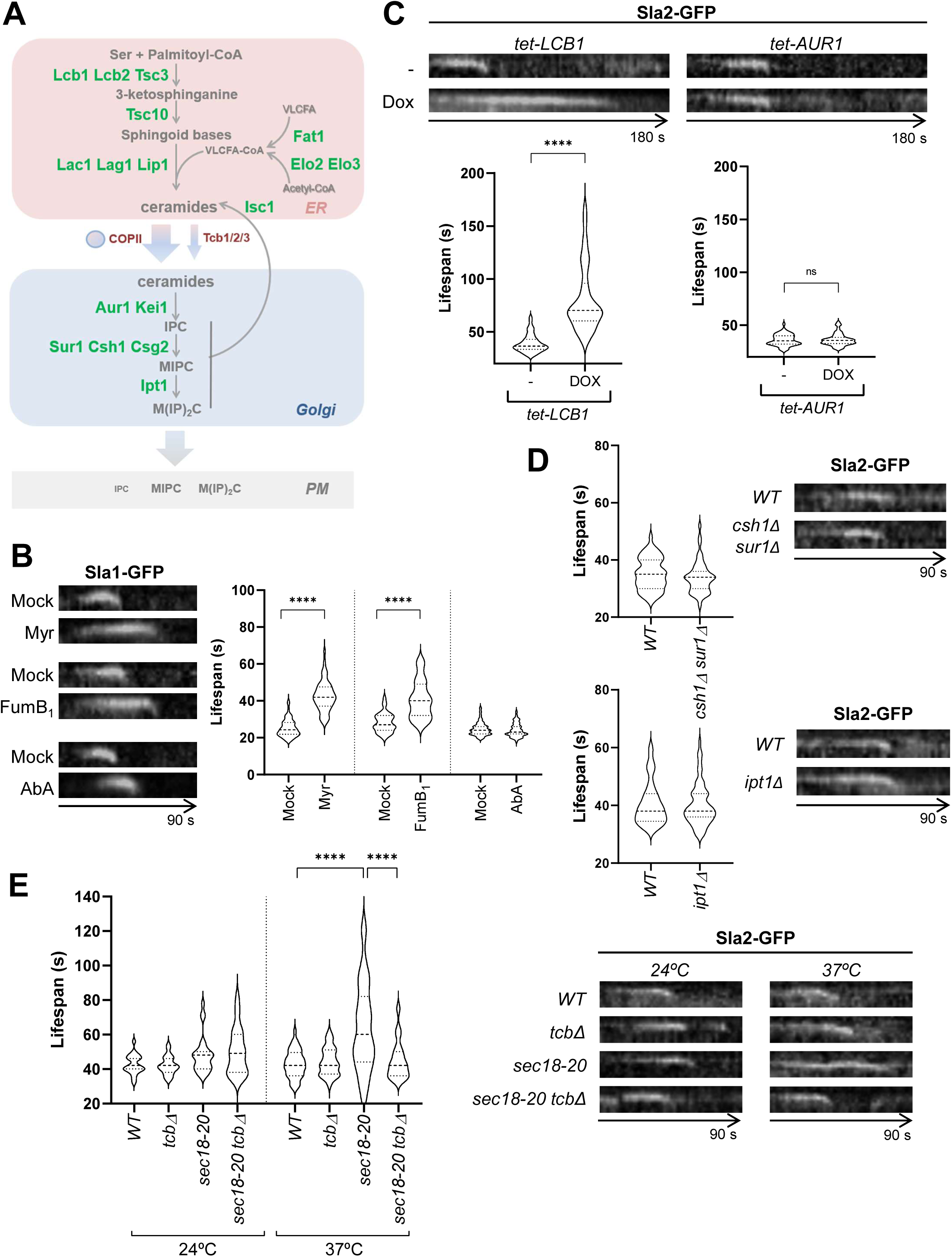
VLCA-Cers but not More Complex Sphingolipids Are Required for Endocytic Uptake. **A.** Scheme depicting the ER and Golgi sphingolipid biosynthesis pathway in yeast. In green are depicted the enzymes involved. **B.** Kymographs of 2 ss time-lapse epi-fluorescence movies of cortical Sla1-GFP patches in wt cells treated with 5 µM Myriocin (Myr), 100 µM Fumonisin B_1_ (FumB_1_) or 0.2 µg/ml Aureobasidin A (AbA), or mock treated; and violin plots of the corresponding Sla1-GFP lifespans. Asterisks indicate statistically significant differences between groups determined by Kruskal-Wallis test with Dunn’s post-hoc correction for multiple comparisons. **C.** Kymographs of 2 ss time-lapse epi-fluorescence movies of cortical Sla2-GFP patches in the indicated strains, treated for 6 hours with or without doxycycline (Dox); and violin plots of the Sla2-GFP cortical patch lifespans. Asterisks indicate statistically significant differences between groups determined by Mann-Whitney test. **D.** Kymographs of 2 ss time-lapse epi-fluorescence movies of cortical Sla2-GFP patches for the indicated strains; and violin plots of the Sla2-GFP cortical patch lifespan for the corresponding strains. Statistical significance was determined by Mann-Whitney test. No differences between groups were found. **E.** Kymographs of 2 s time-lapse epi-fluorescence movies of cortical Sla2-GFP patches for the indicated strains; and violin plots of the Sla2-GFP cortical patch lifespans of in the corresponding strains. Asterisks indicate statistically significant differences between groups determined by Kruskal-Wallis test with Dunn’s post-hoc correction for multiple comparisons. **B to E**. Dashed lines in violin plots indicate the median and dotted lines indicate the interquartile range (25^th^–75^th^ percentiles). n > 50 endocytic patches in more than 20 cells per strain, from a representative experiment.

Finally, we investigated if transport of Cers from the ER to the Golgi might be required for endocytic uptake. In yeast, Cers are mostly transported to the Golgi via vesicular traffic (Funato and Riezman, 2001), but its transport is also contributed by tricalbin-dependent non vesicular-mechanisms (Ikeda et al., 2020). Thus, it has been previously shown that temperature sensitive secretory mutants have residual IPC synthesis at restrictive temperature, that is completely abrogated by combining the secretory mutants with a triple knock out of the yeast tricalbins *TCB1*, *TCB2* and *TCB3* (Fig. 3A) (Ikeda et al., 2020). Consistent with previous observations, we found that a *sec18* conditional mutant exhibited altered endocytic coat dynamics at restrictive temperature, as a result of reduced endocytic cargo load at the PM (Carroll et al., 2012). However, depletion of the tricalbins in this background restored rather than worsened the endocytic defect (Fig 3D), suggesting that Cer accumulation in the ER somehow facilitated endocytic uptake. The data thus suggested that Cers might be delivered to the endocytic sites by a tricalbin-independent non-vesicular transport rather than via the secretory pathway.

### The CerS Interacts with the ERSES Components and the Endocytic Machinery

Absence of VLCFA-Cer would be expected to decrease lipid packing (Castro et al., 2014) and decrease mechanical bending resistance (Kollmitzer et al., 2015); thereby facilitating, rather than slowing down endocytic uptake (Gauthier et al., 2012). Therefore, the data pointed to a specific role of VLCFA-Cers in endocytic internalization, instead of a defect secondarily caused by general changes in the fluidity of the membrane. Indeed, overexpressing of Ole1, the only yeast FA desaturase, known to increase membrane fluidity (Degreif et al., 2017), slightly accelerated the endocytic coat dynamics, as expected (Fig. S3A).

Our observations suggested that VLCA-Cers might directly be transported from the ER to endocytic sites, similar to sterols; and that the CerS might play a direct role in endocytic budding. To investigate if this might be the case, we looked for for physical interactions between the CerS subunits Lac1, Lag1 and Lip1 and both, the ERSES components (Erg2, Erg6, Erg7 and Erg27) and the endocytic machinery, using the yeast two hybrid (Y2H) for an initial screen.

Y2H demonstrated specific interactions of all CerS subunits with Erg6, Erg7 and Erg27 (Fig. 4A & S4A). An immunoprecipitation assay from cER and PM-enriched fractions from cells expressing Lac1-GFP and Erg6-HA, using an anti-GFP antibody, confirmed the interaction of the CerS with the ERSES component (Fig. 4B). Fluorescence microscopy analysis showed localization of Lac1-GFP at the nuclear envelope and the cER as previously shown (Barz and Walter, 1999; Fresques et al., 2015) (Fig. 4C). At the cER tips decorated with Lac1-GFP, the staining coincided with the cortical Erg6-mCherry patches (Fig. 4C), indicating that even though the CerS is not an exclusive component of the ERSES, it is present in these ER subdomains, where it will also encounter Fat1 that will define the length of the acyl chain incorporated in the Cer at these ER sites.

**Figure 4.**
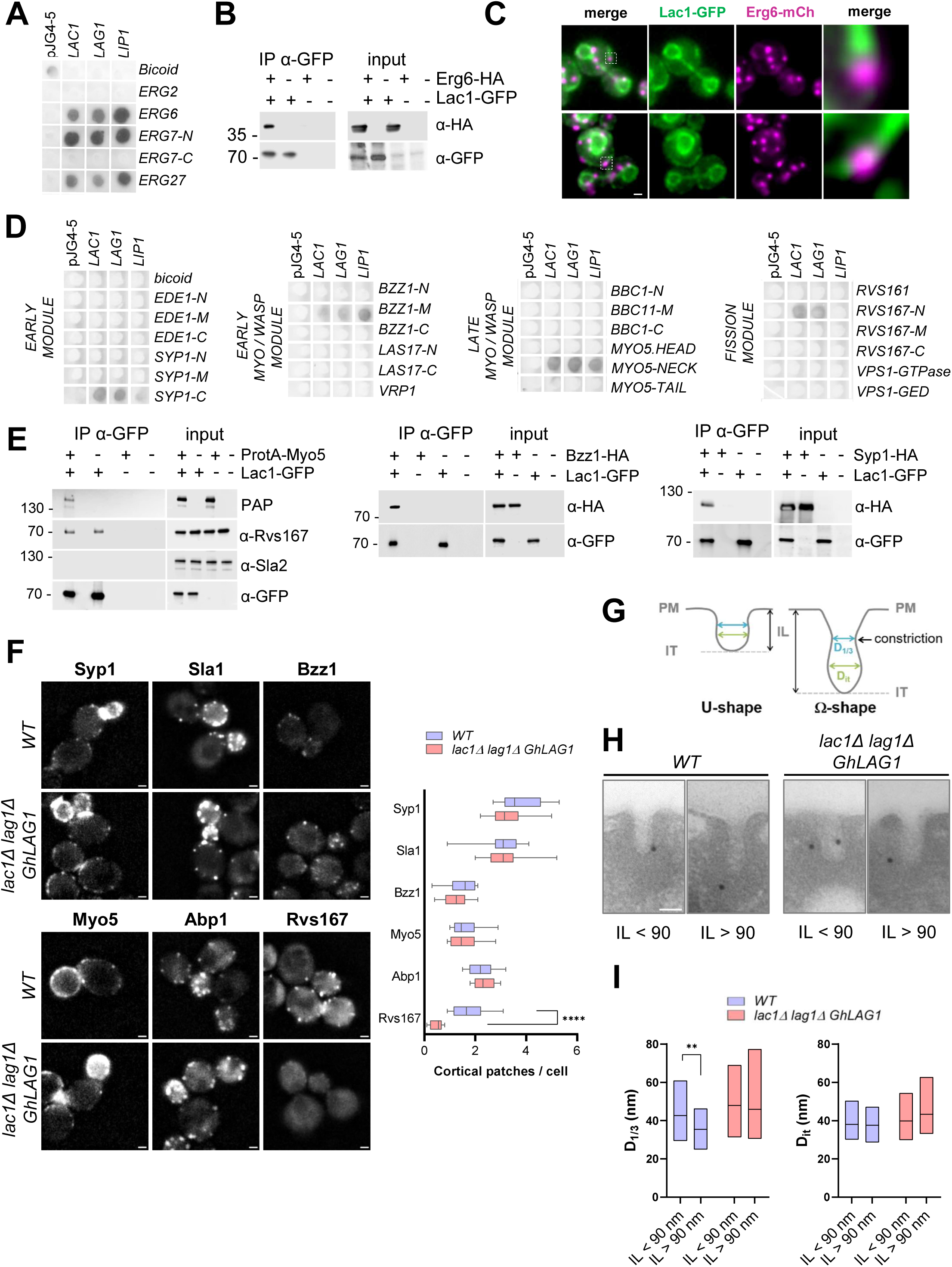
VLCFA-Cers are required for invagination constriction. **A & D.** Y2H assays between the CerS subunits Lac1 Lag1 and Lip1 fused to the B42 transcriptional activator and the indicated Erg proteins (A) or endocytic protein fragments (D) fused to the LexA DNA binding domain, expressed in yeast bearing the β-galactosidase reporter (pSH18-34); grown on X-Gal containing plates. Dark indicates interaction. LexA alone (pJG4-5) or B42 fused to *D. melanogaster* BICOID were used as negative controls. **B.** Immunoblots decorated with anti-GFP (α-GFP) and anti-HA (α-HA) antibodies of anti-GFP agarose immunoprecipitates from non-denaturing protein extracts of yeast expressing the indicated Lac1-GFP (+) or the untagged protein (-); and Erg6-HA (+) or the untagged protein (-). 10 µg of total protein from the extract were loaded as input. **C.** Dual color epi-fluorescent micrographs of live yeast expressing Lac1-GFP and Erg6-mCherry. The individual channels of the same field and the merge are shown. Magnified micrographs of the depicted areas are shown on the right. Bar = 1 µm. **E.** Immunoblots decorated with the indicated antibodies (α-GFP, α-Sla2, α-Rvs167 or α-HA) of anti-GFP agarose immunoprecipitates from non-denaturing protein extracts of yeast expressing the indicated Lac1-GFP (+) or the untagged protein (-); and either ProtA-Myo5 (+) (left panel); Bzz1-HA (+) (middle panel); or Syp1-HA (+) (right panel) or the untagged proteins (-). 10 µg of total protein from the extracts were loaded as input. **F.** Epi-fluorescence micrographs of live wt and *lac1Δ lag1Δ GhLAG1* cells expressing the indicted proteins fused to GFP (Syp1, Sla1, Bzz1, Myo5, Rvs167) or mCherry (Abp1); and box and whiskers plots of the number of the indicated endocytic protein cortical patches in the wt and mutant strains. Boxes represent the 25^th^– 75^th^ percentiles, the line indicates the median, and whiskers represent the 5^th^–95^th^ percentiles. n > 120 cells per strain, from a representative experiment. The number of endocytic patches was only quantified in the mother cells for clarity. Asterisks indicate statistically significant differences between groups determined by Welch’s t-tests. Bar = 1 μm. **G.** Scheme depicting an endocytic invagination and the parameters used to analyze their morphology. PM, plasma membrane; IT, invagination tip; IL, invagination length; D_it_, diameter at the IT; D_1/3_, diameter at 1/3 of the IL. **H.** Electron micrographs of endocytic invaginations of increasing length, decorated with immuno-gold particles against actin in wt and *lac1Δ lag1Δ GhLAG1* cells. Bar = 50 nm. **I.** Low-high plots show the invagination diameter in the indicated location. The line indicates the mean, and bars represent the minimum and maximum values. More than 90 invaginations were analyzed per strain. Statistical significance was determined by two-way ANOVA with Sidak’s multiple comparisons test; asterisks indicate significant differences between groups.

For the endocytic machinery, we focused on proteins that localize at the neck and the PMIJ, where the cER contacts the endocytic invaginations (Encinar Del Dedo et al., 2017). Those proteins included the type I myosin Myo5, the yeast N-WASP and WIP, Las17 and Vrp1; the F-BAR domain proteins Syp1 and Bzz1, the Eps15 homolog Ede1; the yeast N-BAR proteins Rvs161 and Rvs167; and the dynamin Vps1 (Encinar Del Dedo et al., 2017; Idrissi et al., 2012; Idrissi et al., 2008).

Strikingly, numerous interactions of the CerS subunits with the endocytic machinery could be detected. The catalytic CerS subunits showed strongest binding to the Myo5 neck; the µ Homology Domain of Syp1 and the N-BAR domain of Rvs167; whereas Lip1 preferred the intrinsically disordered region of Bzz1 (Fig. 4D and S4A). Immunoprecipitation assays confirmed the interactions of Lac1-GFP with ProtA-Myo5, endogenous Rvs167, Syp1-HA and Bzz1-HA (Fig. 4E). No signal for the endocytic coat component Sla2 could be detected in the Lac1-GFP immunoprecipitates, demonstrating the specificity of the detected interactions (Fig. 4E). The CerS binding site in Myo5 did not overlap with that of Osh3, the Erg proteins or Osh2 (Encinar Del Dedo et al., 2017) (Fig. S4A), further highlighting the scaffolding role of the myosin-I holding the lipid biosynthetic and transport machineries at ER-ECSs.

### VLCA-Cers Facilitate Constriction of Endocytic Invaginations

Constriction of the endocytic invaginations requires the Myo5-ATPase activity and the assembly of N-BAR domain containing proteins Syp1 and Bzz1 at the invagination neck (Idrissi et al., 2012), which is followed by recruitment of Rvs161 and Rvs167 as the invagination further elongates and constricts (Idrissi et al., 2008). The detected interactions between the CerS subunits with Myo5 and the BAR-domain containing proteins suggested that the VLCA-Cers might be involved in constriction of endocytic invaginations.

Consistent with the view that invaginations might not proceed to the latest stage prior to fission in the absence of VLCA-Cers, we found that Rvs167-GFP was not localized in cortical patches in the *lac1Δ lag1Δ GhLAG1* mutant, whereas components of earlier endocytic modules including Syp1, Sla2, Myo5, Bzz1 or Abp1 showed a pattern comparable to wt cells (Fig. 4F). Specific delocalization of Rvs167-GFP was also observed upon treatment with the CerS inhibitor Fumonisin B (Fig. S4B).

To directly investigate if VLCA-Cers are specifically involved in the constriction of endocytic invaginations, we decided to analyze the morphology of the invagination profiles in the *lac1Δ lag1Δ GhLAG1* mutant by quantitative immuno-electron microscopy. *lac1Δ lag1Δ GhLAG1* and wt cells were processed for transmission electron microscopy and endocytic invaginations were labeled with an anti-actin antibody (Idrissi et al., 2008).

Analysis of the morphology of more than 90 endocytic profiles for the wt and CerS mutant strains showed that endocytic invaginations indeed failed to constrict in the absence of VLCA-Cers (Fig. 4G to I). Thus, while the diameter of the invagination tip (D_it_) capped by the endocytic coat (Fig. 4G) remained unchanged throughout the process in both wt and CerS mutant cells (Fig. 4H and I), the diameter measured at one third of the invagination length (D_1/3_) -the site where constriction appears in wt cells (Fernandez-Golbano et al., 2014) (Fig. 4G)-decreased from 44 to 35 nm in wt cells as the tube elongated, whereas it remained constant in the *lac1Δ lag1Δ GhLAG1* mutant (Fig. 4H and I).

### Osh3 and Osh1 are Involved in Cer transport at ER-ECS

Our data indicated that VLCA-Cers play a specific role in constriction of endocytic invaginations. We then speculate, that similar to Osh2, which engages the sterol biosynthetic and endocytic machinery to transfer sterols at ER-ECS (Encinar Del Dedo et al., 2017), other LTPs might hook to the CerS and the endocytic machinery to support VLCFA-Cers transport at ER-ECS. An obvious candidate to test was Osh3, another ER-ECS located ORP that interacts with Myo5 (Encinar Del Dedo et al., 2017) and whose depletion cause Myriocin resistance (Yano et al., 2004) and Aureobasidin A sensitivity (Kajiwara et al., 2014), phenotypes that would be compatible with an *osh3Δ* strain accumulating Cers in the ER.

To check if the Osh3 hydrophobic cavity might be able to accommodate VLCFA-Cers at all, we performed an unbiased coarse-grained molecular dynamics (CG-MD) protocol in which lipids placed away from the protein can spontaneously bind to lipid transfer proteins (Srinivasan et al., 2024). To this end, the Osh3 ORD (aa 668-973) was placed in a box of water with 120 mM NaCl, and a molecule of VLCFA-Cer (d16:0/26:0) was randomly inserted into the bulk solvent at a random position. Then, five independent 1-μs-long unbiased MD simulations were performed, starting from different initial positions of the lipid (Fig. S5A and Video S2). The Osh3 fragment used did not include the lid because the CG-MD simulations cannot detect conformational changes and the lid may block lipid entry as previously observed for Osh4 and Osh6 (Srinivasan et al., 2024). As a control, we also performed MD simulations with phosphatidylinositol 4-phosphate (PI4P) (16:0/18:1), a described Osh3 cargo (Stefan et al., 2011; Tong et al., 2013) (Fig. S5A; Video S3). To assess lipid binding, we used the minimum distance between the protein and the solvation number of the lipid tails (defined as the number of water molecules within 5 Å of the lipid acyl chains) (Fig. S5A). We selected this property since a lipid bound to an external surface of the protein would be surrounded by a larger number of water molecules, as opposed to lipids bound to an hydrophobic cavity. As seen in all MD replicas (Fig. S5A) and in figure 5A, depicting the final binding pose, both VLCFA-Cer and PI4P entered the hydrophobic cavity of Osh3 and remained attached within the simulation time. Solvation analysis during the last 100 ns of the trajectories indicated that the lipid tails of both VLCFA-Cers and PI4P were desolvated at the end of the trajectory, within the range shown for lipid species naturally binding to LTPs, thus confirming binding to the hydrophobic cavity (Srinivasan et al., 2024) (Fig. 5B and S5A). Notably, the disposition of the bound PI4P molecules after the CG-MD protocol closely resembles the experimental structure of the ORD bound to a PI4P molecule (RCSB PDB ID: 4INQ), with the lipid tails inserted into the hydrophobic pocket and the bulky headgroup sticking out of the pocket. The data thus indicated potential promiscuity in the lipids that Osh3 can transfer and suggested that the ORP molecular environment might contribute to dictating its cargo.

**Figure 5.**
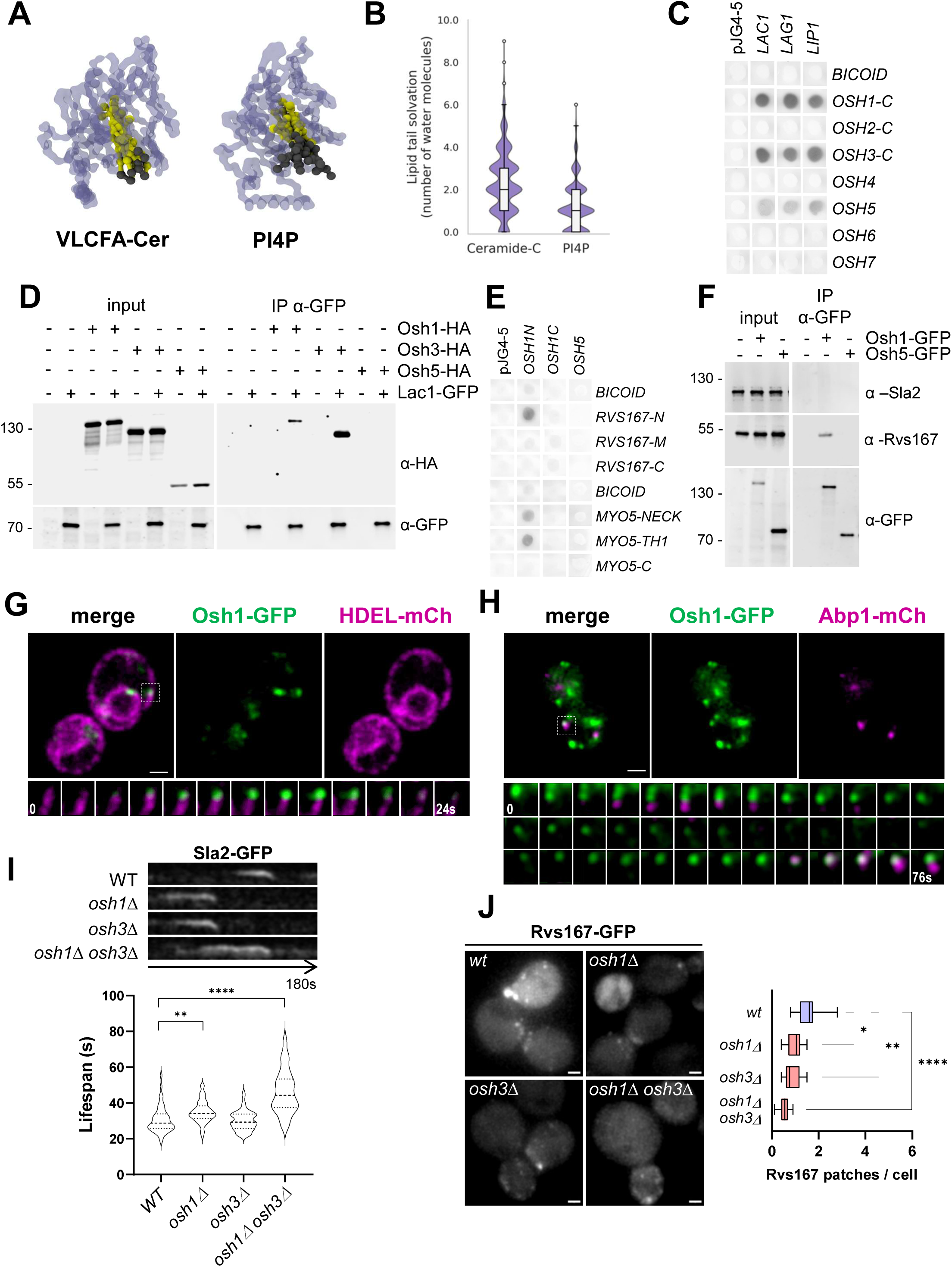
Osh3 and Osh1 Interact with the CerS and the Endocytic Machinery. **A.** Representative final pose of the indicated lipids within the Osh3 ORD after unbiased CG-MD simulations. Structures of the protein were coarse-grained and set up with a lipid randomly placed in a bulk solvent far away from the protein, followed by MD simulations. **B.** Lipid tail solvation in the last 100 ns of the CG-MD trajectories. The two dotted lines indicate the average lipid tail solvation. **C.** Y2H assay between the CerS subunits Lac1 Lag1 and Lip1 fused to the B42 transcriptional activator and the ORDs of the indicated Osh proteins fused to the LexA DNA binding domain, expressed in yeast bearing the β-galactosidase reporter (pSH18-34); grown on X-Gal containing plates. Dark indicates interaction. LexA alone (pJG4-5) or B42 fused to *D. melanogaster* BICOID were used as negative control. **D.** Immunoblots decorated with anti-GFP (α-GFP) or HA (α-HA) of anti-GFP agarose immunoprecipitates from non-denaturing protein extracts of yeast expressing the Lac1-GFP (+) or the untagged protein (-); and either Osh1-HA, Osh3-HA or Osh5-HA (+) or the untagged proteins (-). 10 µg of total protein from the extract were loaded as input. **E.** Y2H assay between the Osh1 N and C-terminal fragments or Osh5 fused to the B42 transcriptional activator, and the fragments of the indicated endocytic proteins fused to the LexA DNA binding domain, expressed in yeast bearing the β-galactosidase reporter (pSH18-34); grown on X-Gal containing plates. Dark indicates interaction. LexA alone (pJG4-5) or B42 fused to *D. melanogaster* BICOID were used as negative control. **F.** Immunoblots decorated with anti-GFP (α-GFP), Rvs167 (α-Rvs167) or Sla2 (α-Sla2) of anti-GFP agarose immunoprecipitates from non-denaturing protein extracts of yeast expressing the indicated Osh proteins fused to GFP (+) or the untagged proteins (-). 10 µg of total protein from the extracts were loaded as input. **G & H.** Dual color confocal-fluorescent micrographs of live yeast expressing Osh1-GFP and HDEL-mCherry (G) or Abp1-mCherry (H). The individual channels and the merged image of the same field are shown. The lower panels show consecutive merged frames from a time-lapse acquired at 2 s intervals, corresponding to the enlarged insets and illustrating the transient colocalization of Osh1-GFP with the rims of cER (G) or the Abp1-mCherry patches (H). Bar = 1 µm. **I.** Kymographs of 2 ss time-lapse epi-fluorescence movies of cortical Sla2-GFP patches for the indicated strains; and violin plots of the Sla2-GFP cortical patch lifespans for the corresponding strains. Asterisks indicate statistically significant differences determined by Kruskal-Wallis test with Dunn’s post-hoc correction for multiple comparisons. **J.** Epi-fluorescence micrographs of the indicated strains expressing Rvs167-GFP; and box and whiskers plots of the number of Rvs167-GFP cortical patches in the wt and mutant strains. Boxes represent the 25^th^–75^th^ percentiles, the line indicates the median, and whiskers represent the 5^th^–95^th^ percentiles. n > 120 cells per strain, from a representative experiment. The number of endocytic patches was only quantified in the mother cells for clarity. Asterisks indicate statistically significant differences between groups determined by Welch’s ANOVA test with Dunnett’s post-hoc correction for multiple comparisons. Bar = 1 μm.

Since we have previously seen that the Osh2 ORD interacts with the Erg machinery and that this interaction is essential to sustain Osh2-dependent sterol extraction from the ERSES (Encinar Del Dedo et al., 2021), we next investigated if the ORD of Osh3 might specifically interact with the CerS. To this end, we tested the ORD domains of all Osh proteins against the CerS subunits Lac1, Lag1 and Lip1 in the Y2H. As shown in figure 5C, the ORD of Osh3, but not Osh2, bound the CerS subunits. Interestingly, Osh1, and to a lesser extent Osh5, also gave positive signals with the CerS, indicating possible functional redundancy. Immunoprecipitation assays confirmed the interaction of Osh3-HA and Osh1-HA, but not Osh5-HA, with Lac1-GFP (Fig. 5D).

To study if Osh1 might work at ER-ECS, we next investigated if it physically interacted with the endocytic machinery. Y2H analysis detected specific interactions of the N-terminal portion of Osh1, but not the full length Osh5, with Myo5 and with Rvs167, pointing to an endocytic function of Osh1 (Fig. 5E). The specific interaction of Osh1-GFP with the endogenous Rvs167 was confirmed by immunoprecipitation (Fig. 5F).

Osh1 was previously reported to localize at the Nuclear-Vacuole Junction (NVJ) and the Golgi (Levine and Munro, 2001). However, the conditions used in these experiments might have overlooked transient interactions with cER and the endocytic sites. 2 s time-lapse live-cell imaging indeed demonstrated that, besides the already reported localization of Osh1 at the NVJ and internal structures, possibly corresponding to the Golgi; small Osh1-GFP patches transiently contacted the rims of the cER (Fig. 5G and movie S4) similar to Osh2 and Osh3 (Encinar Del Dedo et al., 2017). In contrast, Osh5 occupied the surface of the cER facing the PM (Fig. S5B). Remarkably, the cortical Osh1-GFP patches appeared in transient association with the endocytic actin marker Abp1 (Fig. 5H and movie S5), indicating a likely contribution of the LTP at ER-ECSs.

Consistent with a redundant function of Osh1 and Osh3 as Cer transporters at ER-ECSs, double deletion of these ORPs, but not the single mutants, exhibited elongated Sla2-GFP cortical patch life-spans, similar to the *lac1Δ lag1Δ GhLAG1* mutant (Fig. 5I). Also recapitulating the endocytic defects of the strain lacking VLCFA-Cers, the double *osh1Δ osh3Δ* mutant failed to recruit Rvs167 at cortical patches, indicating defects to proceed to the late stages of endocytic budding (Fig. 5J).

To directly test the hypotheses that Cers might be associated with endocytic sites and that Osh1 and Osh3 contribute to Cer export from the ER, we decided to develop a Cer probe to follow these lipid species by fluorescence microscopy in living yeast. To this end, we used the C1 domain of KSR1 (Kinase Suppressor of Ras 1, amino acid 317 to 400, XM_0736985) fused to GFP, which was previously characterized as a Cer probe in mammalian cells (Girik et al., 2024). The cytosolic KSR1-GFP did not demonstrate any specific membrane associated Cer pattern (Fig. S6A), indicating that exposure of the hydroxyl head group of Cers to the cytosolic leaflet of cellular membranes might be residual or not detectable over the cytosolic background under these conditions. In contrast, the same KSR1-GFP probe fused to an ER signal peptide (SP-KSR1-GFP) evidenced a clear ER pattern in co-expression experiments with HDEL-mCherry (Fig. 6A), consistent with the expected localization of Cers in yeast. Treatment of cells with Fumonisin B significantly decreased the SP-KSR1-GFP staining, whereas treatment with Aureobasidin A increased cellular accumulation of the probe (Fig. 6B), demonstrating that the SP-KSR1-GFP also detected yeast Cers. Expression of the SP-KSR1-GFP probe did not significantly altered the dynamics of the cortical endocytic coat under the experimental conditions used (Fig. S6B).

**Figure 6.**
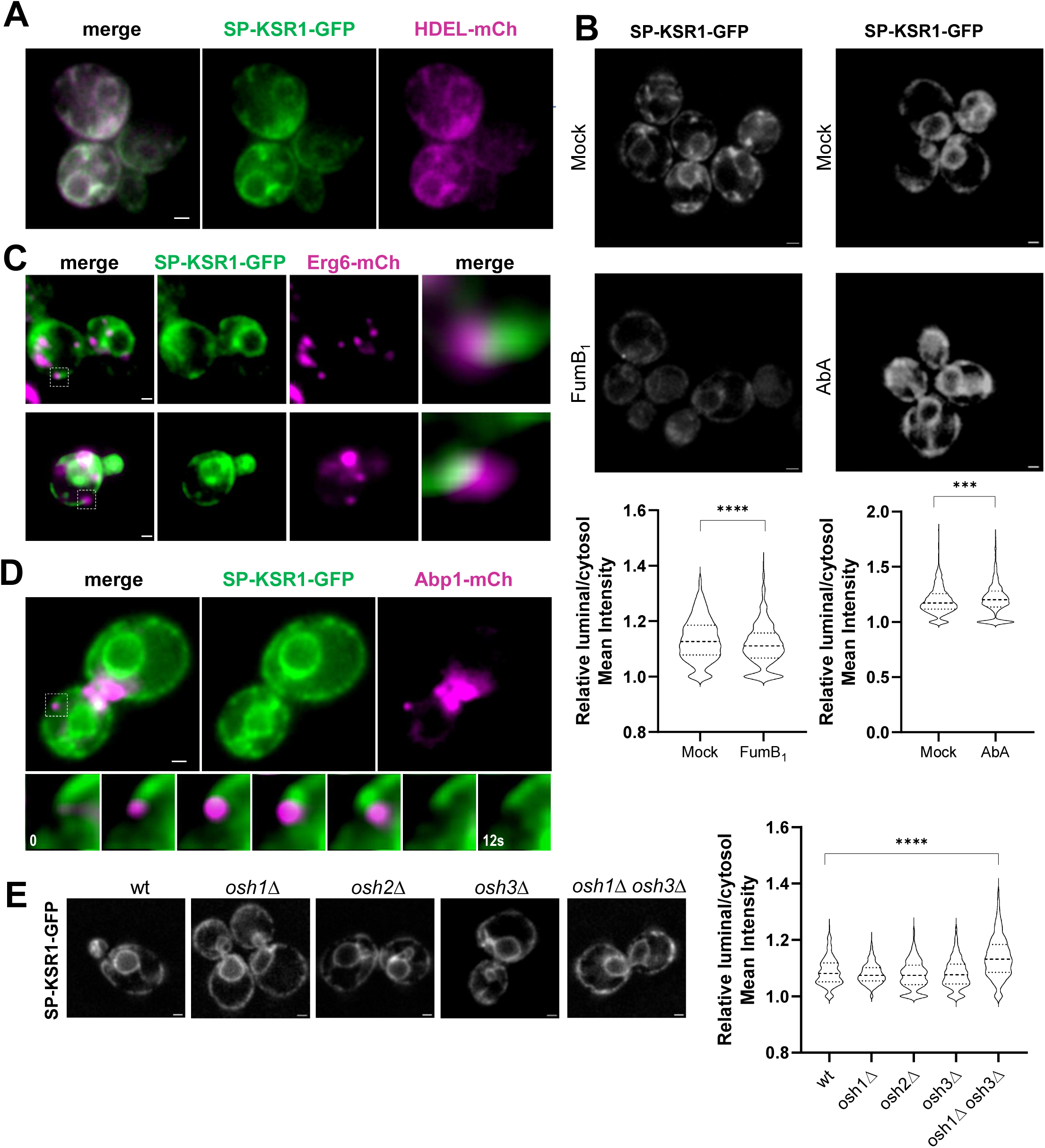
Osh1 and Osh3 contribute to Cer export from the ER. **A.** Dual color epi-fluorescent micrographs of live yeast expressing SP-KSR1-GFP and HDEL-mCherry. The individual channels of the same field and the merge are shown. Bar = 1 µm. **B and E.** Z projections of epi-fluorescence micrographs of yeast expressing SP-KSR1-GFP of either wt cells treated with 100 µM Fumonisin B_1_ (FumB_1_), 0.2 µg/ml Aureobasidin A (AbA), or mock treated (B) or in the indicated strains (E); and violin plot of the SP-KSR1-GFP luminal versus cytosolic normalized mean intensities computed using and automated image analysis pipeline. Asterisks indicate statistically significant differences between groups determined by Mann-Whitney test. n > 400 cells for each strain or experimental condition. Dashed lines in violin plots indicate the median and dotted lines indicate the interquartile range (25th–75th percentiles). Bar = 1 μm. **C.** Dual color epi-fluorescence micrograph of wt live-yeast expressing the SP-KSR1-GFP probe and Erg6-mCherry. The individual channels and the merge of the same fields are shown. The right panels correspond to the magnified insets. Bar = 1 µm. **D.** Dual color live-cell epi-fluorescence micrograph of wt yeast expressing the SP-KSR1-GFP probe and Abp1-mCherry. The individual channels and the merged image of the same field are shown. The lower panel shows consecutive merged frames from a time-lapse acquired at 2 s intervals, corresponding to the enlarged inset. Bar = 1 µm.

Next, we used the probe to investigate possible Cers association with ERSES and endocytic sites. Fluorescence microscopy of yeast cells expressing SP-KSR1-GFP and Erg6-mCherry showed clear coincidence of the fluorescent signals at the cER rims (Fig. 6C). Time-lapse live-cell imaging of yeast expressing SP-KSR1-GFP and Abp1-mCherry also indicated transient association of Cers with endocytic sites (Fig. 6D & movies S6 to 8). Interestingly, the association followed always a similar pattern with the probe accumulating in the interface between Abp1 and the PM, where the PMIJ occurs (Fig. 6D & movies S6 to 8).

Finally, to directly investigate if Osh3 and Osh1 might participate in the extraction of Cers from the ER, we expressed the SP-KSR1-GFP probe in wt and *osh1*Δ and *osh3*Δ single and double mutants and compared their luminal Cer signal to the wt. As minor changes would be expected given that most Cers exit the ER using the secretory pathway (Funato and Riezman, 2001), we developed an automated image analysis pipeline to measure the luminal signal of a large number of cells (https://github.com/MolecularImagingPlatformIBMB/yeast-microscopy-pipeline). As shown in figure 6E, a small but significant increase in the luminal SP-KSR1-GFP signal was observed in the double mutant, as compared with the wt and the single deleted knock outs. No increase in the cER SP-KSR1-GFP luminal signal was detected in *osh2*Δ yeast, which instead accumulates sterols at ERSES (Encinar Del Dedo et al., 2021), suggesting functional specialization (Fig. 6E).

## Discussion

Our data provide strong evidence for a confined and specific role of VLCA-Cers in endocytic uptake, extending well beyond potential nonspecific effects on PM fluidity. In addition to the endocytic defects caused by inhibition of Cer synthesis, substitution of VLCA-Cers with C18-Cers, or depletion of the likely VLCA-Cer transporters at ER–ECS, Osh1 and Osh3; we demonstrate that the CerS physically interacts with several components of the endocytic machinery, pointing to a direct involvement of this enzyme in the process. Furthermore, live-cell imaging using a newly developed probe reveals that Cers accumulate at endocytic sites before the endocytic actin machinery departs into the cytosol, further supporting a temporally defined role for Cers during endocytic membrane remodelling.

A link between Cers and endocytosis was reported in mammalian cells in a variety of contexts, most commonly through the activation or secretion of sphingomyelinases, which hydrolyse sphingomyelin to generate Cer and phosphorylcholine (Tam et al., 2010; (Piacentino et al., 2022; Andrews, 2019).

In artificial giant unilamellar vesicles (GUVs), asymmetric hydrolysis of C16:0 sphingomyelin by exogenously added or microinjected sphingomyelinase drives vectorial budding, leading to the proposal that the cone-shaped geometry of Cers promotes membrane curvature. Together with the propensity of Cers to coalesce and phase-separate from phospholipids, this property can promote membrane vesiculation in the absence of additional molecular machinery (Holopainen et al., 2000). However, this mechanism is largely independent of Cer acyl-chain length and causes massive vesiculation (Holopainen et al., 2000). In contrast, our findings in yeast reveal a spatially and temporally controlled process in which VLCA-Cers are locally transferred at ER–ECS together with sterols and fulfil a confined function during constriction of endocytic invaginations. Importantly, this function is strongly dependent on the acyl-chain length, as C18-Cer poorly substitutes for VLCA-Cer in this context.

Saturated VLCA-Cers possess distinctive biophysical properties. Like other saturated Cers, VLCA-Cers can form gel or crystalline domains that phase-separate from glycerolipids and promote the formation of non-lamellar phases, such as the PMIJ. However, their unusually long acyl chains additionally enable the formation of transbilayer domains and confer strong membrane-deforming properties (Alonso and Goni, 2018; Castro et al., 2014). We speculate that VLCFA-Cers might form a transbilayer lipid boundary at the PMIJ that facilitates generation of membrane tension along the invagination to permit constriction and ultimately fission. What is the exact molecular role of VLCA-Cers in endocytic uptake awaits now definition, but similar to endocytic proteins, they can be considered as an active piece of the membrane deformation machinery in yeast.

Beyond establishing a role for VLCA-Cers in endocytic uptake in yeast, the present work further characterizes the composition and molecular function of the recently identified ERSESs. We demonstrate that, in addition to supporting sterol synthesis and localized transport, ERSESs harbour the machinery required for the synthesis and transport of VLCA-Cers. Our data support a model in which ER sterol and sphingolipid metabolic networks are not organized as linear pathways in which intermediates freely diffuse within the lipid bilayer. Rather, they form nodes where biosynthetic routes physically intersect and coupled to dedicated lipid-transfer systems. This organization may limit the accumulation of functionally disruptive intermediates while facilitating the emergence of transient membrane subdomains with defined biophysical, biochemical and geometrical properties.

A similar physical coupling between lipid biosynthetic enzymes and cellular machineries, mediated by dedicated lipid-transfer proteins (LTPs), may also operate in glycerolipid metabolism. Recruitment of the acyl-CoA synthetase Faa1 to the yeast phagophore, for example, channels C12–C16 fatty acids into local phospholipid synthesis, which is subsequently coupled to Atg2-mediated lipid transport to drive phagophore expansion (Schutter et al., 2020).

The metabolism and transport of sterols and sphingolipids are tightly interconnected by mechanisms that are incompletely understood (Gulati et al., 2010). An imbalance in either lipid invariably results in pathology. In addition to monogenic disorders such as Niemann-Pick diseases, the un-coordinated homeostasis of these lipids has been increasingly implicated in many polygenic disorders such as atherosclerosis and neurodegeneration, including Alzheimeŕs and Huntingtońs diseases (Gulati et al., 2010). ERSESs may represent specialized metabolic and lipid-transfer platforms through which cells coordinate synthesis and transport of different lipids coupled to local membrane remodelling, offering a potential entry point for deciphering the still poorly understood functional interplay between sterols and sphingolipids in health and disease.

## Authors contributions

ISG and LH, led the project and performed the FM on the sphingolipid pathway and on the Osh proteins and the ceramide probe, respectively; with assistance of IG. ISG performed the QIEM experiments. LH performed the Y2H assays and immunoprecipitations with contributions from OT, IG, SICX and MIG. ER guided confocal microscopy and develop IA-and macro-based pipelines for wide-field image quantification. DA performed the MD experiments. ISG, LH, MIG, IG, OT, SICX constructed strains and plasmids. MIG supervised, secured funding and wrote the article.

## Acknowledgements

This work was supported by the MINECO (PID2023-150264NB-I00) and PIF_24_00374 to L. H. We thank T. Garcia for assistance with FM, P. Carvalho (Oxford University), K. Funato (Hirochima University), H. Riezman (University of Geneve) and O. Gallego (Universitat Pompeu Fabra) for sending strains or plasmids. DA acknowledges the Centro Informático Científico de Andalucía (CICA) for providing the computational resources for the CG-MD simulations.

## STAR methods

### Key Resource Table

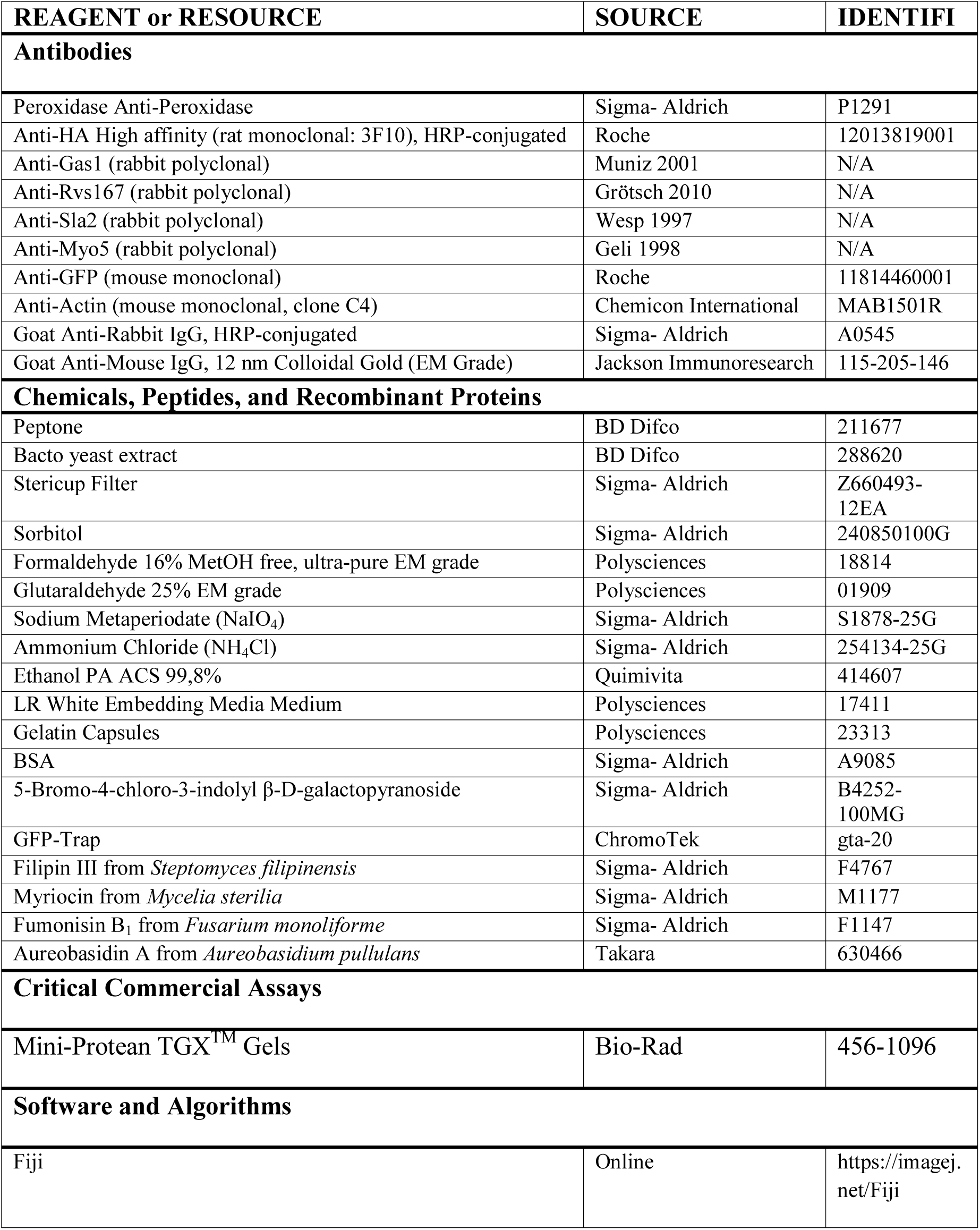
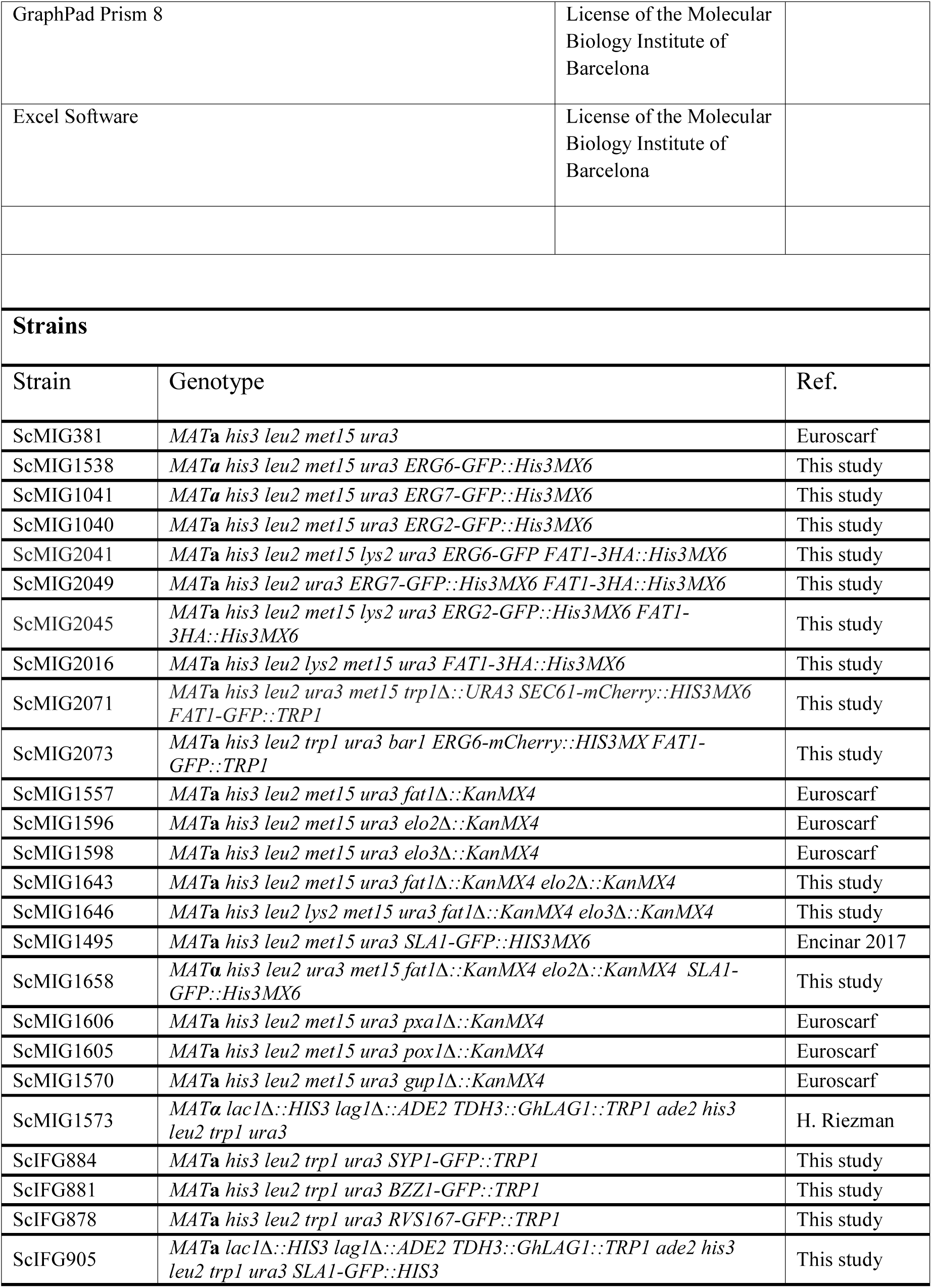
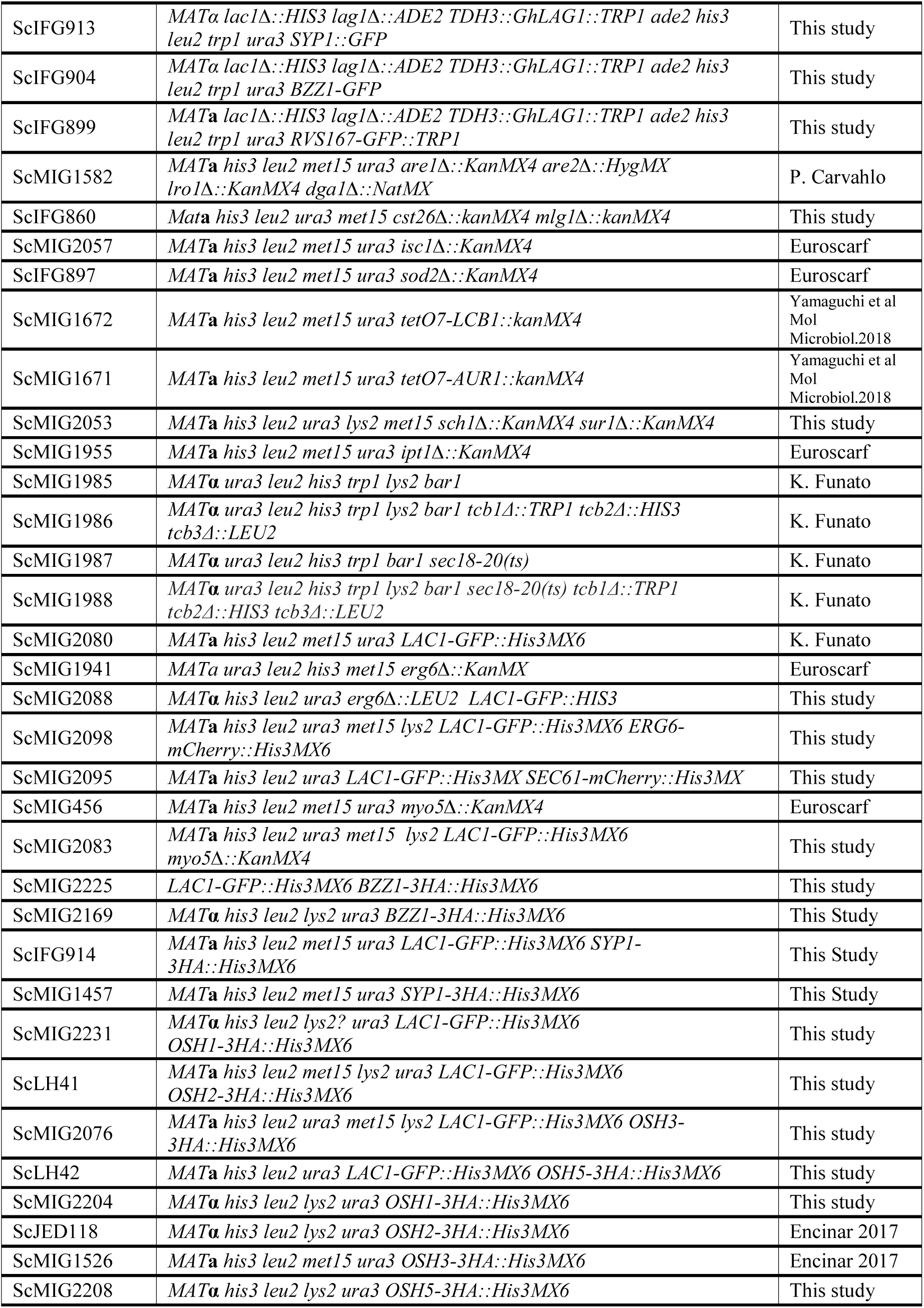
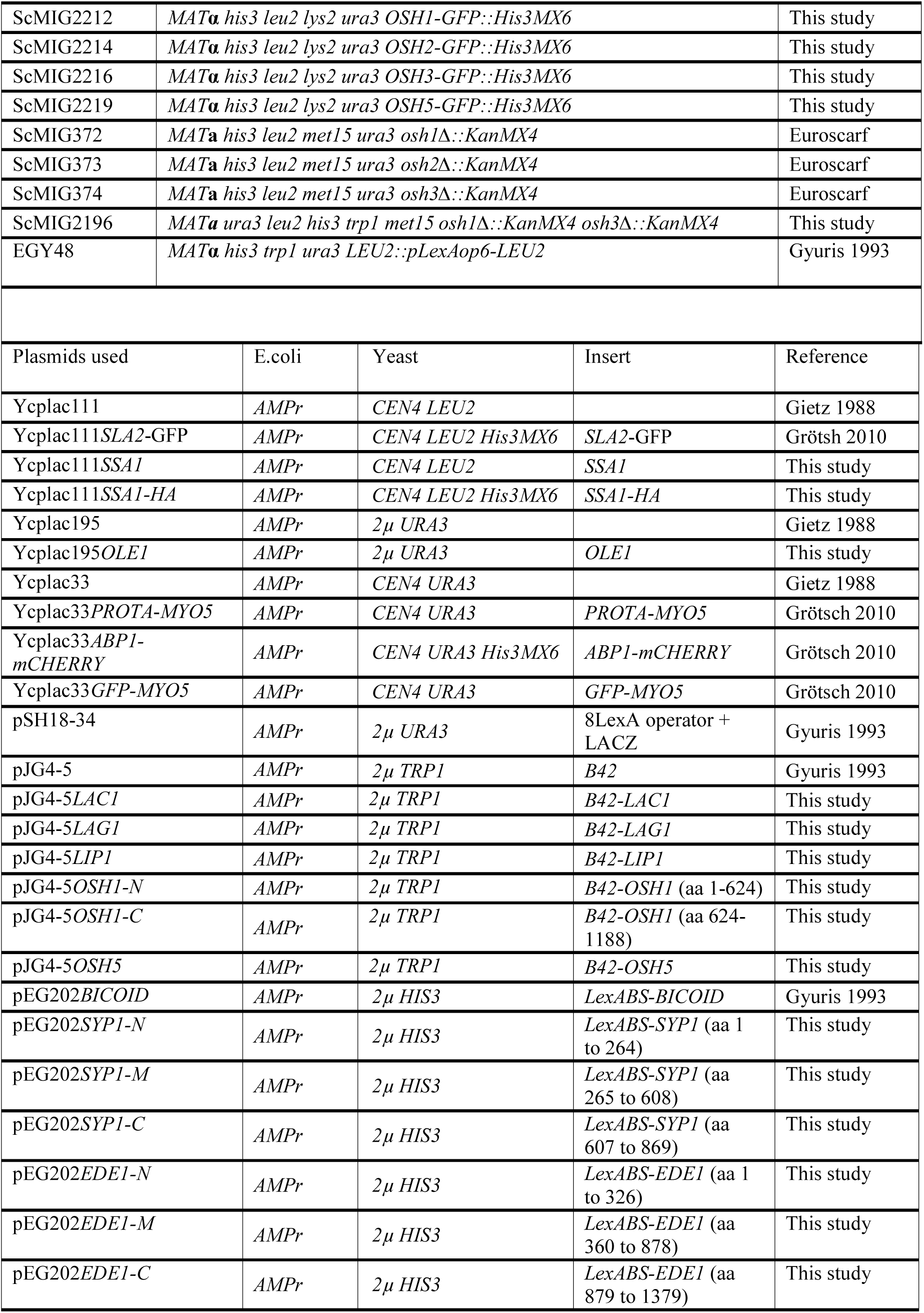
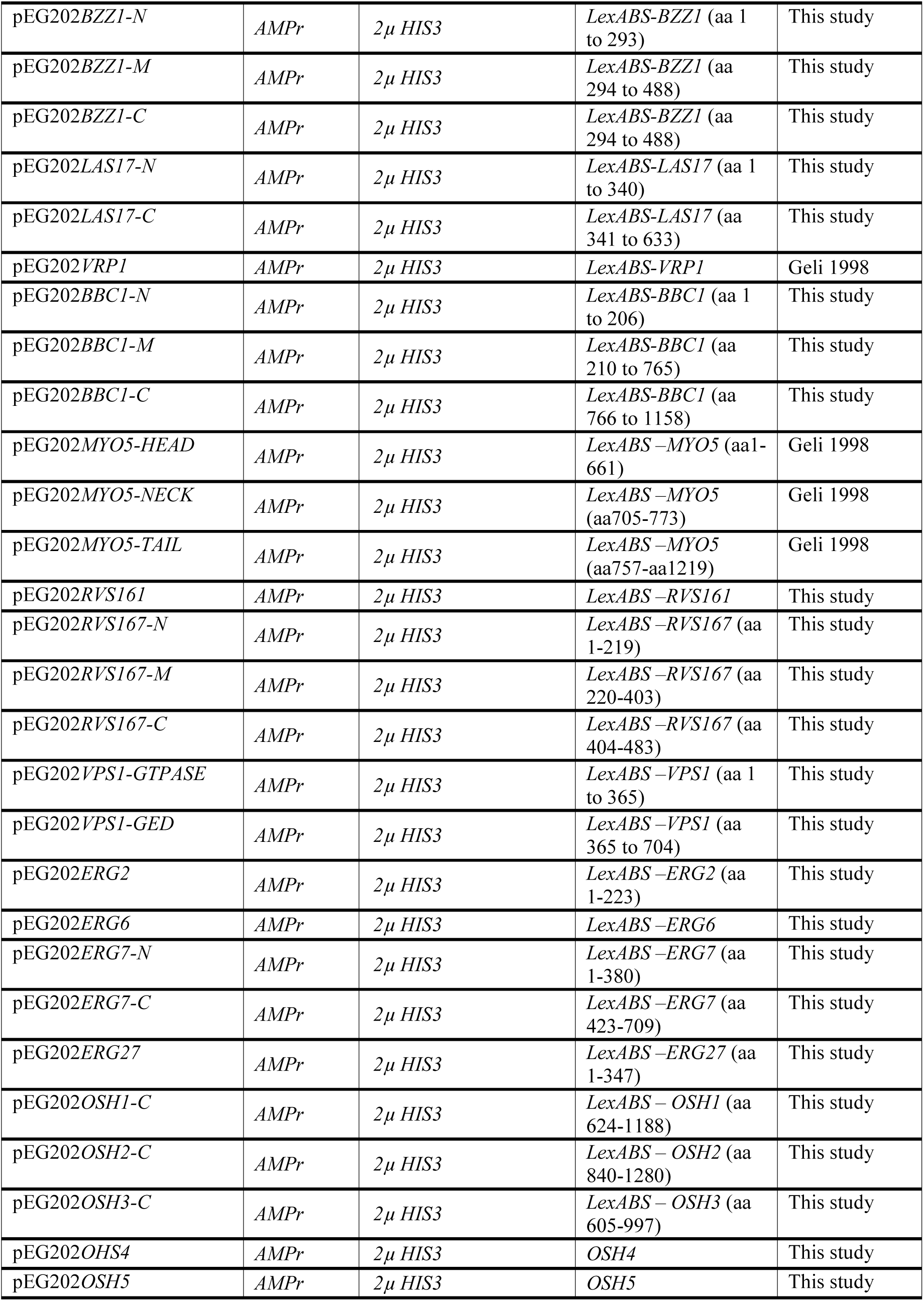
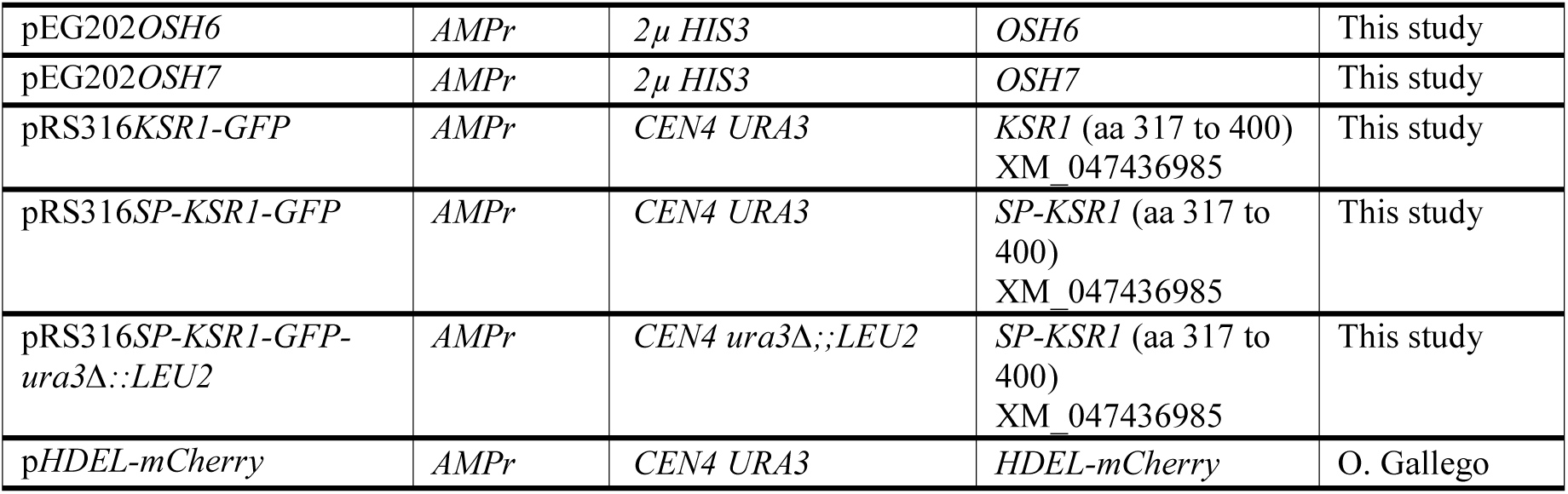

| REAGENT or RESOURCE | SOURCE | IDENTIFI |  |  |
|---|---|---|---|---|
| **Antibodies** |  |  |  |  |
| Peroxidase Anti-Peroxidase | Sigma- Aldrich | P1291 |  |  |
| Anti-HA High affinity (rat monoclonal: 3F10), HRP-conjugated | Roche | 12013819001 |  |  |
| Anti-Gas1 (rabbit polyclonal) | Muniz 2001 | N/A |  |  |
| Anti-Rvs167 (rabbit polyclonal) | Grötsch 2010 | N/A |  |  |
| Anti-Sla2 (rabbit polyclonal) | Wesp 1997 | N/A |  |  |
| Anti-Myo5 (rabbit polyclonal) | Geli 1998 | N/A |  |  |
| Anti-GFP (mouse monoclonal) | Roche | 11814460001 |  |  |
| Anti-Actin (mouse monoclonal, clone C4) | Chemicon International | MAB1501R |  |  |
| Goat Anti-Rabbit IgG, HRP-conjugated | Sigma- Aldrich | A0545 |  |  |
| Goat Anti-Mouse IgG, 12 nm Colloidal Gold (EM Grade) | Jackson Immunoresearch | 115-205-146 |  |  |
| **Chemicals, Peptides, and Recombinant Proteins** |  |  |  |  |
| Peptone | BD Difco | 211677 |  |  |
| Bacto yeast extract | BD Difco | 288620 |  |  |
| Stericup Filter | Sigma- Aldrich | Z660493-12EA |  |  |
| Sorbitol | Sigma- Aldrich | 240850100G |  |  |
| Formaldehyde 16% MetOH free, ultra-pure EM grade | Polysciences | 18814 |  |  |
| Glutaraldehyde 25% EM grade | Polysciences | 01909 |  |  |
| Sodium Metaperiodate (NaIO<sub>4</sub>) | Sigma- Aldrich | S1878-25G |  |  |
| Ammonium Chloride (NH<sub>4</sub>Cl) | Sigma- Aldrich | 254134-25G |  |  |
| Ethanol PA ACS 99,8% | Quimivita | 414607 |  |  |
| LR White Embedding Media Medium | Polysciences | 17411 |  |  |
| Gelatin Capsules | Polysciences | 23313 |  |  |
| BSA | Sigma- Aldrich | A9085 |  |  |
| 5-Bromo-4-chloro-3-indolyl β-D-galactopyranoside | Sigma- Aldrich | B4252-100MG |  |  |
| GFP-Trap | ChromoTek | gta-20 |  |  |
| Filipin III from *Steptomyces filipinensis* | Sigma- Aldrich | F4767 |  |  |
| Myriocin from *Mycelia sterilia* | Sigma- Aldrich | M1177 |  |  |
| Fumonisin B<sub>1</sub> from *Fusarium monoliforme* | Sigma- Aldrich | F1147 |  |  |
| Aureobasidin A from *Aureobasidium pullulans* | Takara | 630466 |  |  |
| **Critical Commercial Assays** |  |  |  |  |
| Mini-Protean TGX<sup>TM</sup> Gels | Bio-Rad | 456-1096 |  |  |
| **Software and Algorithms** |  |  |  |  |
| Fiji | Online | https://imagej.net/Fiji |  |  |
| GraphPad Prism 8 | License of the Molecular Biology Institute of Barcelona |  |  |  |
| Excel Software | License of the Molecular Biology Institute of Barcelona |  |  |  |
| **Strains** |  |  |  |  |
| Strain | Genotype | Ref. |  |  |
| ScMIG381 | *MAT**a** his3 leu2 met15 ura3* | Euroscarf |  |  |
| ScMIG1538 | *MAT**a** his3 leu2 met15 ura3 ERG6-GFP::His3MX6* | This study |  |  |
| ScMIG1041 | *MAT**a** his3 leu2 met15 ura3 ERG7-GFP::His3MX6* | This study |  |  |
| ScMIG1040 | *MAT**a** his3 leu2 met15 ura3 ERG2-GFP::His3MX6* | This study |  |  |
| ScMIG2041 | *MAT**a** his3 leu2 met15 lys2 ura3 ERG6-GFP FAT1-3HA::His3MX6* | This study |  |  |
| ScMIG2049 | *MAT**a** his3 leu2 ura3 ERG7-GFP::His3MX6 FAT1-3HA::His3MX6* | This study |  |  |
| ScMIG2045 | *MAT**a** his3 leu2 met15 lys2 ura3 ERG2-GFP::His3MX6 FAT1-3HA::His3MX6* | This study |  |  |
| ScMIG2016 | *MAT**a** his3 leu2 lys2 met15 ura3 FAT1-3HA::His3MX6* | This study |  |  |
| ScMIG2071 | *MAT**a** his3 leu2 ura3 met15 trp1Δ::URA3 SEC61-mCherry::HIS3MX6 FAT1-GFP::TRP1* | This study |  |  |
| ScMIG2073 | *MAT**a** his3 leu2 trp1 ura3 bar1 ERG6-mCherry::HIS3MX FAT1-GFP::TRP1* | This study |  |  |
| ScMIG1557 | *MAT**a** his3 leu2 met15 ura3 fat1Δ::KanMX4* | Euroscarf |  |  |
| ScMIG1596 | *MAT**a** his3 leu2 met15 ura3 elo2Δ::KanMX4* | Euroscarf |  |  |
| ScMIG1598 | *MAT**a** his3 leu2 met15 ura3 elo3Δ::KanMX4* | Euroscarf |  |  |
| ScMIG1643 | *MAT**a** his3 leu2 met15 ura3 fat1Δ::KanMX4 elo2Δ::KanMX4* | This study |  |  |
| ScMIG1646 | *MAT**a** his3 leu2 lys2 met15 ura3 fat1Δ::KanMX4 elo3Δ::KanMX4* | This study |  |  |
| ScMIG1495 | *MAT**a** his3 leu2 met15 ura3 SLA1-GFP::HIS3MX6* | Encinar 2017 |  |  |
| ScMIG1658 | *MAT**α** his3 leu2 ura3 met15 fat1Δ::KanMX4 elo2Δ::KanMX4 SLA1-GFP::His3MX6* | This study |  |  |
| ScMIG1606 | *MAT**a** his3 leu2 met15 ura3 pxa1Δ::KanMX4* | Euroscarf |  |  |
| ScMIG1605 | *MAT**a** his3 leu2 met15 ura3 pox1Δ::KanMX4* | Euroscarf |  |  |
| ScMIG1570 | *MAT**a** his3 leu2 met15 ura3 gup1Δ::KanMX4* | Euroscarf |  |  |
| ScMIG1573 | *MAT**α** lac1Δ::HIS3 lag1Δ::ADE2 TDH3::GhLAG1::TRP1 ade2 his3 leu2 trp1 ura3* | H. Riezman |  |  |
| ScIFG884 | *MAT**a** his3 leu2 trp1 ura3 SYP1-GFP::TRP1* | This study |  |  |
| ScIFG881 | *MAT**a** his3 leu2 trp1 ura3 BZZ1-GFP::TRP1* | This study |  |  |
| ScIFG878 | *MAT**a** his3 leu2 trp1 ura3 RVS167-GFP::TRP1* | This study |  |  |
| ScIFG905 | *MAT**a** lac1Δ::HIS3 lag1Δ::ADE2 TDH3::GhLAG1::TRP1 ade2 his3 leu2 trp1 ura3 SLA1-GFP::HIS3* | This study |  |  |
| ScIFG913 | *MATα lac1Δ::HIS3 lag1Δ::ADE2 TDH3::GhLAG1::TRP1 ade2 his3 leu2 trp1 ura3 SYP1::GFP* | This study |  |  |
| ScIFG904 | *MATα lac1Δ::HIS3 lag1Δ::ADE2 TDH3::GhLAG1::TRP1 ade2 his3 leu2 trp1 ura3 BZZ1-GFP* | This study |  |  |
| ScIFG899 | *MAT**a** lac1Δ::HIS3 lag1Δ::ADE2 TDH3::GhLAG1::TRP1 ade2 his3 leu2 trp1 ura3 RVS167-GFP::TRP1* | This study |  |  |
| ScMIG1582 | *MAT**a** his3 leu2 met15 ura3 are1Δ::KanMX4 are2Δ::HygMX lro1Δ::KanMX4 dga1Δ::NatMX* | P. Carvahlo |  |  |
| ScIFG860 | *Mat**a** his3 leu2 ura3 met15 cst26Δ::kanMX4 mlg1Δ::kanMX4* | This study |  |  |
| ScMIG2057 | *MAT**a** his3 leu2 met15 ura3 isc1Δ::KanMX4* | Euroscarf |  |  |
| ScIFG897 | *MAT**a** his3 leu2 met15 ura3 sod2Δ::KanMX4* | Euroscarf |  |  |
| ScMIG1672 | *MAT**a** his3 leu2 met15 ura3 tetO7-LCB1::kanMX4* | Yamaguchi et al Mol Microbiol.2018 |  |  |
| ScMIG1671 | *MAT**a** his3 leu2 met15 ura3 tetO7-AUR1::kanMX4* | Yamaguchi et al Mol Microbiol.2018 |  |  |
| ScMIG2053 | *MAT**a** his3 leu2 ura3 lys2 met15 sch1Δ::KanMX4 sur1Δ::KanMX4* | This study |  |  |
| ScMIG1955 | *MAT**a** his3 leu2 met15 ura3 ipt1Δ::KanMX4* | Euroscarf |  |  |
| ScMIG1985 | *MAT**α** ura3 leu2 his3 trp1 lys2 bar1* | K. Funato |  |  |
| ScMIG1986 | *MAT**α** ura3 leu2 his3 trp1 lys2 bar1 tcb1Δ::TRP1 tcb2Δ::HIS3 tcb3Δ::LEU2* | K. Funato |  |  |
| ScMIG1987 | *MAT**α** ura3 leu2 his3 trp1 bar1 sec18-20(ts)* | K. Funato |  |  |
| ScMIG1988 | *MAT**α** ura3 leu2 his3 trp1 lys2 bar1 sec18-20(ts) tcb1Δ::TRP1 tcb2Δ::HIS3 tcb3Δ::LEU2* | K. Funato |  |  |
| ScMIG2080 | *MAT**a** his3 leu2 met15 ura3 LAC1-GFP::His3MX6* | K. Funato |  |  |
| ScMIG1941 | *MATa ura3 leu2 his3 met15 erg6Δ::KanMX* | Euroscarf |  |  |
| ScMIG2088 | *MAT**α** his3 leu2 ura3 erg6Δ::LEU2 LAC1-GFP::HIS3* | This study |  |  |
| ScMIG2098 | *MAT**a** his3 leu2 ura3 met15 lys2 LAC1-GFP::His3MX6 ERG6-mCherry::His3MX6* | This study |  |  |
| ScMIG2095 | *MAT**a** his3 leu2 ura3 LAC1-GFP::His3MX SEC61-mCherry::His3MX* | This study |  |  |
| ScMIG456 | *MAT**a** his3 leu2 met15 ura3 myo5Δ::KanMX4* | Euroscarf |  |  |
| ScMIG2083 | *MAT**a** his3 leu2 ura3 met15 lys2 LAC1-GFP::His3MX6 myo5Δ::KanMX4* | This study |  |  |
| ScMIG2225 | *LAC1-GFP::His3MX6 BZZ1-3HA::His3MX6* | This study |  |  |
| ScMIG2169 | *MAT**α** his3 leu2 lys2 ura3 BZZ1-3HA::His3MX6* | This Study |  |  |
| ScIFG914 | *MAT**a** his3 leu2 met15 ura3 LAC1-GFP::His3MX6 SYP1-3HA::His3MX6* | This Study |  |  |
| ScMIG1457 | *MAT**a** his3 leu2 met15 ura3 SYP1-3HA::His3MX6* | This Study |  |  |
| ScMIG2231 | *MAT**α** his3 leu2 lys2? ura3 LAC1-GFP::His3MX6 OSH1-3HA::His3MX6* | This study |  |  |
| ScLH41 | *MAT**a** his3 leu2 met15 lys2 ura3 LAC1-GFP::His3MX6 OSH2-3HA::His3MX6* | This study |  |  |
| ScMIG2076 | *MAT**a** his3 leu2 ura3 met15 lys2 LAC1-GFP::His3MX6 OSH3-3HA::His3MX6* | This study |  |  |
| ScLH42 | *MAT**a** his3 leu2 ura3 LAC1-GFP::His3MX6 OSH5-3HA::His3MX6* | This study |  |  |
| ScMIG2204 | *MAT**α** his3 leu2 lys2 ura3 OSH1-3HA::His3MX6* | This study |  |  |
| ScJED118 | *MAT**α** his3 leu2 lys2 ura3 OSH2-3HA::His3MX6* | Encinar 2017 |  |  |
| ScMIG1526 | *MAT**a** his3 leu2 met15 ura3 OSH3-3HA::His3MX6* | Encinar 2017 |  |  |
| ScMIG2208 | *MAT**α** his3 leu2 lys2 ura3 OSH5-3HA::His3MX6* | This study |  |  |
| ScMIG2212 | *MAT**α** his3 leu2 lys2 ura3 OSH1-GFP::His3MX6* | This study |  |  |
| ScMIG2214 | *MAT**α** his3 leu2 lys2 ura3 OSH2-GFP::His3MX6* | This study |  |  |
| ScMIG2216 | *MAT**α** his3 leu2 lys2 ura3 OSH3-GFP::His3MX6* | This study |  |  |
| ScMIG2219 | *MAT**α** his3 leu2 lys2 ura3 OSH5-GFP::His3MX6* | This study |  |  |
| ScMIG372 | *MAT**a** his3 leu2 met15 ura3 osh1Δ::KanMX4* | Euroscarf |  |  |
| ScMIG373 | *MAT**a** his3 leu2 met15 ura3 osh2Δ::KanMX4* | Euroscarf |  |  |
| ScMIG374 | *MAT**a** his3 leu2 met15 ura3 osh3Δ::KanMX4* | Euroscarf |  |  |
| ScMIG2196 | *MAT**a** ura3 leu2 his3 trp1 met15 osh1Δ::KanMX4 osh3Δ::KanMX4* | This study |  |  |
| EGY48 | *MAT**α** his3 trp1 ura3 LEU2::pLexAop6-LEU2* | Gyuris 1993 |  |  |
| Plasmids used | E.coli | Yeast | Insert | Reference |
| Ycplac111 | *AMPr* | *CEN4 LEU2* |  | Gietz 1988 |
| Ycplac111*SLA2*-GFP | *AMPr* | *CEN4 LEU2 His3MX6* | *SLA2*-GFP | Grötsh 2010 |
| Ycplac111*SSA1* | *AMPr* | *CEN4 LEU2* | *SSA1* | This study |
| Ycplac111*SSA1-HA* | *AMPr* | *CEN4 LEU2 His3MX6* | *SSA1-HA* | This study |
| Ycplac195 | *AMPr* | *2μ URA3* |  | Gietz 1988 |
| Ycplac195*OLE1* | *AMPr* | *2μ URA3* | *OLE1* | This study |
| Ycplac33 | *AMPr* | *CEN4 URA3* |  | Gietz 1988 |
| Ycplac33*PROTA-MYO5* | *AMPr* | *CEN4 URA3* | *PROTA-MYO5* | Grötsch 2010 |
| Ycplac33*ABP1-mCHERRY* | *AMPr* | *CEN4 URA3 His3MX6* | *ABP1-mCHERRY* | Grötsch 2010 |
| Ycplac33*GFP-MYO5* | *AMPr* | *CEN4 URA3* | *GFP-MYO5* | Grötsch 2010 |
| pSH18-34 | *AMPr* | *2μ URA3* | 8LexA operator + LACZ | Gyuris 1993 |
| pJG4-5 | *AMPr* | *2μ TRP1* | *B42* | Gyuris 1993 |
| pJG4-5*LAC1* | *AMPr* | *2μ TRP1* | *B42-LAC1* | This study |
| pJG4-5*LAG1* | *AMPr* | *2μ TRP1* | *B42-LAG1* | This study |
| pJG4-5*LIP1* | *AMPr* | *2μ TRP1* | *B42-LIP1* | This study |
| pJG4-5*OSH1-N* | *AMPr* | *2μ TRP1* | *B42-OSH1* (aa 1-624) | This study |
| pJG4-5*OSH1-C* | *AMPr* | *2μ TRP1* | *B42-OSH1* (aa 624-1188) | This study |
| pJG4-5*OSH5* | *AMPr* | *2μ TRP1* | *B42-OSH5* | This study |
| pEG202*BICOID* | *AMPr* | *2μ HIS3* | *LexABS-BICOID* | Gyuris 1993 |
| pEG202*SYP1-N* | *AMPr* | *2μ HIS3* | *LexABS-SYP1* (aa 1 to 264) | This study |
| pEG202*SYP1-M* | *AMPr* | *2μ HIS3* | *LexABS-SYP1* (aa 265 to 608) | This study |
| pEG202*SYP1-C* | *AMPr* | *2μ HIS3* | *LexABS-SYP1* (aa 607 to 869) | This study |
| pEG202*EDE1-N* | *AMPr* | *2μ HIS3* | *LexABS-EDE1* (aa 1 to 326) | This study |
| pEG202*EDE1-M* | *AMPr* | *2μ HIS3* | *LexABS-EDE1* (aa 360 to 878) | This study |
| pEG202*EDE1-C* | *AMPr* | *2μ HIS3* | *LexABS-EDE1* (aa 879 to 1379) | This study |
| pEG202*BZZ1-N* | *AMPr* | *2μ HIS3* | *LexABS-BZZ1* (aa 1 to 293) | This study |
| pEG202*BZZ1-M* | *AMPr* | *2μ HIS3* | *LexABS-BZZ1* (aa 294 to 488) | This study |
| pEG202*BZZ1-C* | *AMPr* | *2μ HIS3* | *LexABS-BZZ1* (aa 294 to 488) | This study |
| pEG202*LAS17-N* | *AMPr* | *2μ HIS3* | *LexABS-LAS17* (aa 1 to 340) | This study |
| pEG202*LAS17-C* | *AMPr* | *2μ HIS3* | *LexABS-LAS17* (aa 341 to 633) | This study |
| pEG202*VRP1* | *AMPr* | *2μ HIS3* | *LexABS-VRP1* | Geli 1998 |
| pEG202*BBC1-N* | *AMPr* | *2μ HIS3* | *LexABS-BBC1* (aa 1 to 206) | This study |
| pEG202*BBC1-M* | *AMPr* | *2μ HIS3* | *LexABS-BBC1* (aa 210 to 765) | This study |
| pEG202*BBC1-C* | *AMPr* | *2μ HIS3* | *LexABS-BBC1* (aa 766 to 1158) | This study |
| pEG202*MYO5-HEAD* | *AMPr* | *2μ HIS3* | *LexABS –MYO5* (aa1-661) | Geli 1998 |
| pEG202*MYO5-NECK* | *AMPr* | *2μ HIS3* | *LexABS –MYO5* (aa705-773) | Geli 1998 |
| pEG202*MYO5-TAIL* | *AMPr* | *2μ HIS3* | *LexABS –MYO5* (aa757-aa1219) | Geli 1998 |
| pEG202*RVS161* | *AMPr* | *2μ HIS3* | *LexABS –RVS161* | This study |
| pEG202*RVS167-N* | *AMPr* | *2μ HIS3* | *LexABS –RVS167* (aa 1-219) | This study |
| pEG202*RVS167-M* | *AMPr* | *2μ HIS3* | *LexABS –RVS167* (aa 220-403) | This study |
| pEG202*RVS167-C* | *AMPr* | *2μ HIS3* | *LexABS –RVS167* (aa 404-483) | This study |
| pEG202*VPS1-GTPASE* | *AMPr* | *2μ HIS3* | *LexABS –VPS1* (aa 1 to 365) | This study |
| pEG202*VPS1-GED* | *AMPr* | *2μ HIS3* | *LexABS –VPS1* (aa 365 to 704) | This study |
| pEG202*ERG2* | *AMPr* | *2μ HIS3* | *LexABS –ERG2* (aa 1-223) | This study |
| pEG202*ERG6* | *AMPr* | *2μ HIS3* | *LexABS –ERG6* | This study |
| pEG202*ERG7-N* | *AMPr* | *2μ HIS3* | *LexABS –ERG7* (aa 1-380) | This study |
| pEG202*ERG7-C* | *AMPr* | *2μ HIS3* | *LexABS –ERG7* (aa 423-709) | This study |
| pEG202*ERG27* | *AMPr* | *2μ HIS3* | *LexABS –ERG27* (aa 1-347) | This study |
| pEG202*OSH1-C* | *AMPr* | *2μ HIS3* | *LexABS – OSH1* (aa 624-1188) | This study |
| pEG202*OSH2-C* | *AMPr* | *2μ HIS3* | *LexABS – OSH2* (aa 840-1280) | This study |
| pEG202*OSH3-C* | *AMPr* | *2μ HIS3* | *LexABS – OSH3* (aa 605-997) | This study |
| pEG202*OHS4* | *AMPr* | *2μ HIS3* | *OSH4* | This study |
| pEG202*OSH5* | *AMPr* | *2μ HIS3* | *OSH5* | This study |
| pEG202*OSH6* | *AMPr* | *2μ HIS3* | *OSH6* | This study |
| pEG202*OSH7* | *AMPr* | *2μ HIS3* | *OSH7* | This study |
| pRS316*KSR1-GFP* | *AMPr* | *CEN4 URA3* | *KSR1* (aa 317 to 400)<br>XM_047436985 | This study |
| pRS316*SP-KSR1-GFP* | *AMPr* | *CEN4 URA3* | *SP-KSR1* (aa 317 to 400)<br>XM_047436985 | This study |
| pRS316*SP-KSR1-GFP-ura3Δ::LEU2* | *AMPr* | *CEN4 ura3Δ;;LEU2* | *SP-KSR1* (aa 317 to 400)<br>XM_047436985 | This study |
| p*HDEL-mCherry* | *AMPr* | *CEN4 URA3* | *HDEL-mCherry* | O. Gallego |

### CONTACT FOR REAGENT AND RESOURCE SHARING

Further information and requests for reagents may be directed to María Isabel Geli.

### Strains and Growth Conditions

Yeast strains used are listed in the Key Resource Table. GFP, mCherry, YFP and HA tags were fused at the C-terminus of each protein by homologous recombination in the genome as described previously (Wach et al., 1997). Genome edited strains behaved as wts. Strains without plasmids were grown in complete yeast peptone dextrose (YPD) and strains with plasmids were selected on synthetic dextrose complete (SDC) lacking the appropriate nutrient (Dulic et al., 1991) at 25°C. Transformation of yeast was accomplished by the lithium acetate method (Ito et al., 1983).

### DNA Techniques and Plasmid Construction

DNA manipulations were performed as described (Sambrook et al., 1989). Enzymes for molecular biology were obtained from New England Biolabs. PCRs were performed with a Vent polymerase (New England Biolabs) and a TRIO-thermoblock (Biometra GmbH). Plasmids used in this study are listed in the Key Resource Table. Oligonucleotides used for plasmids construction are listed in the Table S1. Information about the construction strategies is available upon request.

### SDS–PAGE, Immunoblots and Antibodies

SDS–PAGE was performed as described (Laemmli, 1970) using pre-casted Mini-PROTEAN^®^ TGX 4-20% acrylamide gels (Bio-Rad). For immunoblot, nitrocellulose membranes (Protran 0.45 NC, Cytiva) were probed with anti-HA (Anti-HA-Peroxidase High Affinity 3F10, Roche) or the indicated antibodies, followed by a peroxidase-conjugated secondary goat anti-rabbit (for anti-Gas1, Rvs167, Sla2 or Myo5) IgG antibodies (Sigma-Aldrich). ProtA-Myo5 was detected with a Peroxidase conjugated, peroxidase complex (PAP) (Sigma-Aldrich). Protein transfer, blotting and chemiluminescence detection were performed using standard procedures. Detection of proteins was performed using the ECL kit (Cytiva). Co-immunoprecipitated proteins were detected with an Odyssey M Imaging System (LICORbio).

### Subcellular Fractionation and Immunoprecipitations

For the anti-GFP pull downs from P13000 extracts, cells were glass bead-lysed in LB (25 mM Tris, 5 mM EDTA, pH 8.5) in the presence of protease inhibitors (PIs) (1 mM PMSF, 5 µg/ml Leupeptine, 2,5 µg/ml Antipain, 1 µg/ml Pepstatin and 1 µg/ml Aprotinin). Unbroken cells were eliminated at 700 g for 15 min at 4°C. The supernatant was diluted with the same volume of BB (10 mM Tris, 0.2 mM EDTA, 0.2 mM DTT, pH 7.5) containing PIs. The post 700 g supernatants were spin at 13000 g for 20 min, the pellet was recovered and resuspended in 1 ml of IP buffer (50 mM Tris, 150 mM NaCl, 5 mM EDTA, 1% Triton X-100, pH 7.5). The extract was adjusted to 1 mg of protein/ml, incubated 30 minutes at 4°C in the presence of gluthathione-Sepharose (Cytiva) to eliminate unspecific binding and spin at 700 g for 15 min. The supernatant was then incubated with either 30 µl of 50% GFP-Trap (Chromotek) for 1 h at 4°C. Beads were washed with IP, and subsequently with IP buffer without Triton X-100 and boiled in 40 µl SDS-PAGE sample buffer.

### Two Hybrid Assay

The Interaction Trap two-hybrid system was used (Gyuris et al., 1993). Plasmids pEG202, pJG4-5, pSH18-34 and the strain EGY48 were kindly provided by Dr. Brent. To measure β-galactosidase activity, EGY48 cells bearing the lexAop-lacZ reporter plasmid pSH18-34 were co-transformed with the appropriate pEG202 and pJG4-5 derived plasmids (see below) and streaked out on X-Gal (5-bromo-4-chloro-3-indolyl-β-D-galactopyranoside) containing SD/Gal/Raf-His-Trp-Ura plates (0.67 % yeast nitrogen base (Difco), 7 g/l Na_2_HPO_4_, 3 g/l NaH_2_PO_4_, 2% galactose, 1% raffinose, 40 mg/ml leucine, 80 mg/l X-Gal (Sigma-Aldrich), 2% agar, pH 7). Pictures were taken after 2 days of growth at 24°C.

### Fluorescence Microscopy

Cells were grown to mid-log phase, harvested, resuspended in a small volume of SDC medium and imaged as wet mounts of 2% agarose pads prepared in the same medium and at the same temperature used for culture growth. Time-lapse movies were acquired with a 2 s interval. Unless otherwise mentioned, samples were observed on a THUNDER Imager 3D live cell wide-field microscope (Leica Microsystems), equipped with a 63×/1.2 NA water immersion objective, a DFC9000 sCMOS camera, and a Spectra-X Light Engine. GFP was excited and detected using BP475/28 and BP515/40 filters, respectively. mCherry was excited and detected using BP575/35 and BP641/78 filters. Hardware-driven Adaptive Focus Control (AFC) was used to maintain focus over time. The system was controlled using LAS X (Leica Application Suite X) software.

Cells were treated for 2 h with 5 µg/ml Filipin, 100 µM Fumonisin B_1_, 5 µM Myriocin and 0.2 µg/ml Aureobasidin A when indicated. Mock-treated cells received the corresponding vehicle: dimethyl sulfoxide (DMSO) for Filipin and Fumonisin B_1_, methanol for Myriocin, and ethanol for Aureobasidin A, at the same final concentration (v/v) as in the drug-treated samples. Treatment with DOX was extended to 6 hours with 10 µg/ml and 0.2 µg/ml for the *tet-AUR1* and *tet-LCB1* strains, respectively.

Confocal live-cell imaging was performed on a STELLARIS 8 FALCON microscope (Leica Microsystems) equipped with a 63x/1.2 NA water immersion objective. To minimize photobleaching and phototoxicity during live acquisition, images were acquired in photon counting mode, which allowed for a drastic reduction in excitation laser power. To compensate for the low signal-to-noise ratio typical of low expression levels, line accumulation (8 to 16 lines) was applied. Two-channel time-lapse movies (xyt) were acquired by line sequential scanning at a 2 s interval, using a pixel size of 0.07 × 0.07 µm. GFP and mCherry conjugates were excited using the 489 nm and 587 nm lines of a white light laser (WLL), and their emissions collected using sensitive HyD detectors in the 494-586 and 597-818 nm detection windows respectively.

### Image Analysis and Ceramide Quantification

To quantify the subcellular accumulation of the ceramide probe, an automated image analysis pipeline was developed using Fiji/ImageJ (Schindelin et al., 2012) and Python. First, three-channel 3D hyperstacks (GFP, mCherry-HDEL, and BF) were preprocessed and merged into a single 2D composite image to maximize structural information for segmentation. Fluorescence channels were processed using background subtraction, median filtering, and maximum intensity projections, and were subsequently combined with edge-detected median projections of the brightfield channel. Whole-cell segmentation masks were then generated from these synthetic composite images using a custom Cellpose model (Stringer et al., 2021) trained on the cyto3 base model, implemented via a custom Python script.

Subsequently, a custom Fiji macro was used to evaluate the fluorescence ratio between accumulated ceramide structures and the diffuse cytosolic signal. Segmented cell ROIs were filtered by minimum area and solidity to exclude debris and poorly segmented cells. To computationally isolate the concentrated ceramide signal, a global high-intensity mask was generated from the GFP channel using a filtering sequence that included Gaussian blur, rolling-ball background subtraction, gamma adjustment, Tubeness filtering, and bandpass filtering, followed by Moments thresholding. For each individual cell, the accumulated ceramide ROI was defined by the logical intersection (AND) of the cell boundary and the global high-intensity mask, while the diffuse cytosolic ROI was obtained by subtracting this accumulated signal from the whole cell area (XOR). Finally, the mean fluorescence intensity of the GFP signal was measured for both compartments to calculate the accumulated-to-cytosolic ceramide ratio.

All the open-source scripts and the custom cellpose model, along with a description of the pipeline can be found at https://github.com/MolecularImagingPlatformIBMB/yeast-microscopy-pipeline.

### Quantitative Immuno-Electron Microscopy

Immunoelectron microscopy was performed as described in (Idrissi et al., 2008). Briefly, cells were grown in yeast extract peptone dextrose medium (YPD) to 4-5 x 10^6^ cells/ml, harvested over a disposable Stericup 0.22 μm filter unit, leaving 5 ml of media. 25 ml of 1.2x fixative solution was immediately added to obtain final concentrations of 0.04 M KPO4 pH 6.6; 0.6 M sorbitol, 4% formaldehyde (Polysciences), 0.4% glutaraldehyde (Polysciences), 1 mM MgCl_2_, 0.5 mM EGTA (Ethylene-Glycol-bis(2-aminoethylether)-N,N,N’,N’-tetraacetic acid), 10 mM NaF and 10 mM NaN_3_. Cells were then transferred to a 50 ml Falcon and fixation was continued overnight at 4 °C with rolling. Subsequent steps (metaperiodate and ammonium chloride treatments, dehydration, infiltration, embedding and sectioning) were performed as described (Mulholland et al., 1994) without further modifications. For immuno-labeling, ultrathin sections were incubated for 15 min in blocking buffer (10 mM KPO4 pH 7.5, 150 mM NaCl, 2% BSA (bovine serum albumin), 0.05% Tween-20), transferred to a 25 μl drop of primary antibody in blocking buffer for 3 hours, and washed over 30 minutes in washing buffer (10 mM KPO4 Buffer pH 7.5, 150 mM NaCl, 0.05% Tween-20). After blocking for 15 min, grids were incubated with the corresponding gold-conjugated secondary antibody for 60 min and washed first for 30 minutes in washing buffer and 30 minutes in washing buffer without Tween-20. All steps were performed at room temperature. Grids were then washed in double distilled water, fixed for 20 minutes in 8% Glutaraldehyde, and post-stained with uranyl acetate (2% in water) over 30 minutes and lead citrate for 30 s as described (Reynolds, 1963). 50 ug/ml of anti-actin monoclonal antibody (C4; Chemicon International) was used as primary antibody. 12 nm gold-conjugated anti-mouse (Jackson ImmunoResearch Laboratories) was used as a secondary antibody (diluted 1:50). Ultrathin sections were examined using a Jeol 1010 transmission electron microscope at 50KV accelerating voltage. Micrographs of the yeast PM invaginations were acquired at 100.000x magnification with a MegaView III CCD camera and an image acquisition software analySIS (Soft Imaging System, Münster, Germany). Adjustments of image size, brightness and contrast were performed on ImageJ

### Quantification and Statistical Analysis

Quantifications and Kymographs were performed with ImageJ (Rasband, WS, NIH, Bethesda, Maryland, USA, 1997-2016). Statistical tests were selected based on the number of groups compared, the distribution of the data, and the experimental design. For comparisons between two groups, Mann-Whitney test was used when data did not follow a normal distribution; Welch’s t-test was used when data were normally distributed but variances were unequal. For comparisons among three or more groups, Kruskal-Wallis test with Dunn’s post-hoc correction for multiple comparisons was used for non-normally distributed data. For the analysis of invagination diameter across invagination length and strains, two-way ANOVA with Sidak’s multiple comparisons test was used. For categorical data Chi-square test was used to compare proportions between groups. Statistical analyses were performed using GraphPad Prism 8. Asterisks denote statistical significance as follows: *p<0.05, **p<0.01, ***p<0.001, ****p<0.0001; ns, not significant."

### Molecular Dynamics Simulations Methods: CG-MD simulations

Molecular dynamics simulations were performed with the GROMACS (v 2023.3) package (Van Der Spoel et al., 2005), and the MARTINI 3 force field (Souza et al., 2021). Visual Molecular Dynamics (VMD) (Humphrey et al., 1996) was used for visualization and the renders.

The atomistic structure of Osh3 was obtained from the X-Ray structure from RCSD PDB with ID: 4IC4 (Tong et al., 2013) and converted to coarse-grain (CG) using the Martinize script (de Jong et al., 2013). Only residues 669 to 974 of the protein were considered for the simulations, as the presence of the lid in related proteins prevents the binding of lipids (Srinivasan et al., 2024). An additional elastic network with a force constant of 1,000 kJ·mol^-1^·nm^-2^ was used to restrain the secondary structure of the protein, with a 0 nm elastic bond lower cutoff and an upper cutoff of 0.8 nm.

The CG model of Osh3 was then placed in the center of a cubic box, with at least 2.0 nm distance between the protein and the box edge. One lipid molecule was randomly placed in the bulk solvent in each of the ten replicas, without any distance restrictions. The system was then solvated, considering a Van der Waals distance of 0.21 nm, neutralized and ionized with 0.12 M NaCl. Each replica was simulated for 1 µs.

Initial equilibration was carried out by performing energy minimization using the steepest descent algorithm, followed by a short MD run of 125 ps. Production runs were performed at 310 K using a velocity-rescale thermostat (Bussi et al., 2007), with separate temperature coupling for protein and nonprotein particles. The md integrator was used for the production runs, with a time step of 25 fs. The Parrinello–Rahman barostat (Parrinello and Rahman, 1981) was used to maintain a pressure of 1 bar, along with an isotropic pressure coupling scheme and a nstpcouple parameter set to 10. The Coulombic terms were calculated using the reaction field, with an epsilon (dielectric constant) of 15 and a cutoff distance of 1.1 nm. A cutoff scheme was applied for the Van der Waals terms, using a cutoff distance of 1.1 nm, and the Verlet cutoff scheme was used for the potential shift (van Eerden et al., 2017). The non-bonded interactions were calculated by generating a pair-list using the Verlet scheme with a buffer tolerance of 0.005.

## Analysis

A tcl script was used to compute the solvation number of the lipids, counting the number of water molecules within 5.0 Å of the lipid tail beads (headgroup and backbone beads were excluded) for each trajectory frame, for each replica. The tcl script is freely available at the repository https://github.com/danialv4/Unbiased_simulations_characterize_lipid_binding/ To compute the minimum distance between the lipid and the protein, the GROMACS tool mindist was employed. It has been already proved that a simultaneous low minimum distance and low solvation number is indicative of lipid binding to the lipid-transfer-protein (Srinivasan et al., 2024).

## Supplementary Movies Legends

**Movie S1: Cortical Fat1-GFP Patches Are Hot Spots for Assembly of Endocytic Actin.** 2 s epi-fluorescence movies of Fat1-GFP cortical patches from yeast also expressing the endocytic actin marker Abp1-mCherry.

**Movie S2: Entry of VLCFA-Cer into the Osh3 ORD.** 1 µs movie of a representative CG-MD simulation of a VLCFA-Cer entering the hydrophobic cavity of Osh3. The movie runs at 30 frames per s.

**Video S3: Entry of PI4P into the Osh3 ORD.** 1 µs movie of a representative CG-MD simulation of PI4P entering the hydrophobic cavity of Osh3. The movie runs at 30 frames per s.

**Video S4: Osh1-GFP Patches Are Recruited to the cER Rims.** 2 s time-lapse confocal-fluorescence movies of Osh1-GFP patches from yeast also expressing Sec61-mCherry.

**Video S5: Osh1-GFP Cortical Patches Coincide with Endocytic Actin.** 2 s time-lapse confocal-fluorescence movies of Osh1-GFP cortical patches from yeast also expressing Abp1-mCherry.

**Video S6 to 8: Dynamics of Cers at Endocytic Sites.** 2 s epi-fluorescence movies of Abp1-mCherry cortical patches in yeast also expressing SP-KSR1-GFP.

## Supplementary data

### Supplementary Figure Legends

**Figure S1.**
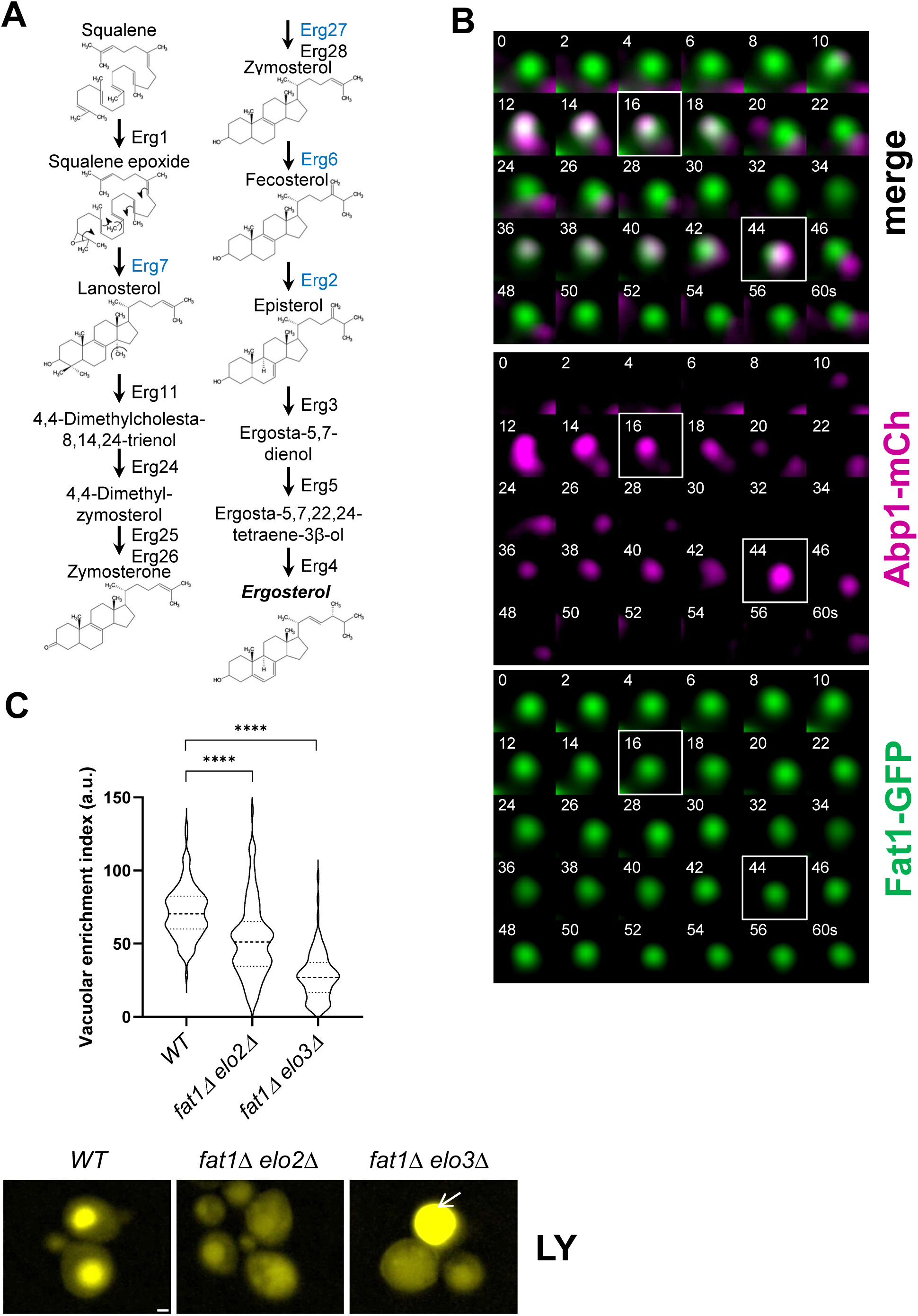
VLCFA-CoAs Are Required for Endocytic Uptake. **A.** Scheme depicting the late ergosterol biosynthetic pathway. Enzymes found in ERSES are depicted in blue **B.** Consecutive merged frames from a dual color epi-fluorescent time-lapse movie acquired at 2 seconds intervals illustrating the transient colocalization of Fat1-GFP with Abp1-mCherry patches. **C.** Epi-fluorescence micrographs of the indicated yeast strains exposed to 4 mg/ml of Lucifer Yellow for one hour; and violin plots of the relative vacuolar intensities (mean intensity at the vacuole-mean intensity at the cytosol). Asterisks indicate significant differences between groups determined by Kruskal-Wallis test with Dunn’s post-hoc correction for multiple comparisons. Dashed lines in violin plots indicate the median and dotted lines indicate the interquartile range (25^th^–75^th^ percentiles). n > 150 cells per strain, from a representative experiment. Bar = 1 μm.

**Figure S2.**
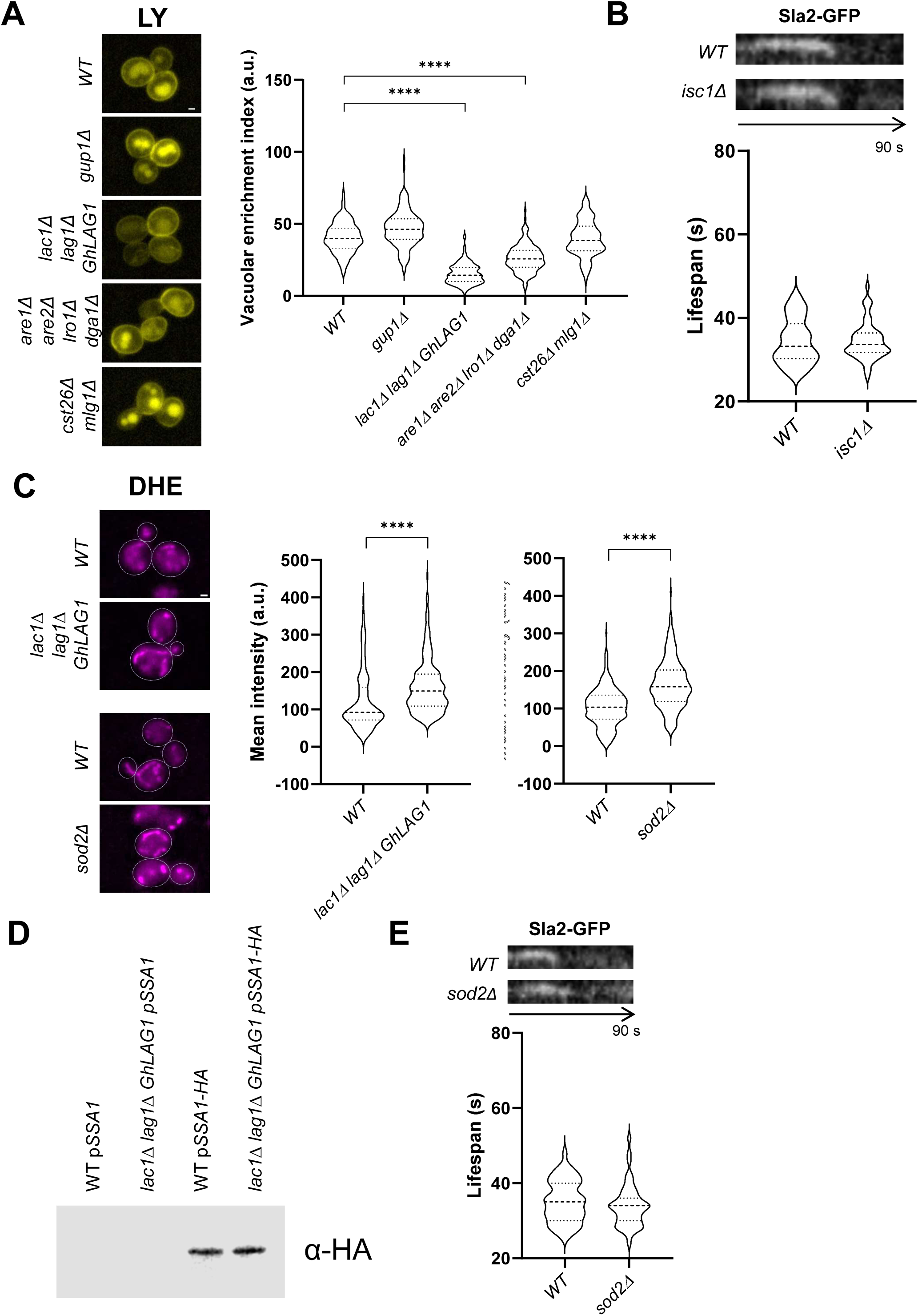
Cer Synthesis, but not other VLCFA-CoAs Anabolic or Catabolic Pathways, Is Required for Endocytic Uptake. **A.** Epi-fluorescence micrographs of the indicated yeast strains exposed to 4 mg/ml of Lucifer yellow for one hour; and violin plots of the relative vacuolar intensities (mean intensity at the vacuole-mean intensity at the cytosol) for the corresponding strains. Asterisks indicate statistically significant differences between groups determined by Kruskal-Wallis test with Dunn’s post-hoc correction for multiple comparisons. **B.** Kymographs of 2 ss time-lapse epi-fluorescence movies of cortical Sla2-GFP patches for the indicated strains; and violin plots of the Sla2-GFP cortical patch lifespans in the corresponding strains. Statistical significance was determined by Mann-Whitney test. No differences between groups were found. **C.** Epi-fluorescence micrographs of the indicated yeast strains stained with DHE for 30 minutes; and violinplots of the mean cellular DHE fluorescence intensity (a. u.) of the corresponding strains. Asterisks indicate statistical significant differences between groups determined by Mann-Whitney test. Bar = 1 μm. Asterisks indicate statistically significant differences between groups determined by Mann-Whitney test. **D.** Immunoblot decorated with an anti-HA (α-HA) antibody of protein extracts from wt or *lac1Δ lag1Δ GhLAG1* strains either bearing a centromeric plasmid expressing *SSA1* (p*SSA1*) or HA-tagged SSA1 (p*SSA1-HA*). 1 µg of total protein was loaded per lane. **E.** Kymographs of 2 s time-lapse epi-fluorescence movies of cortical Sla2-GFP patches for the indicated strains; and violin plots of the Sla2-GFP cortical patch lifespans for the corresponding strains. Statistical significance was determined by Mann-Whitney test. No differences between groups were found. **A, B, C & E.** Dashed lines in the violin plots indicate the median and dotted lines indicate the interquartile range (25^th^–75^th^ percentiles). n > 150 cells (A & C) and n > 50 patches from more than 20 cells (B & E) were analyzed per strain; from a representative experiment

**Figure S3.**
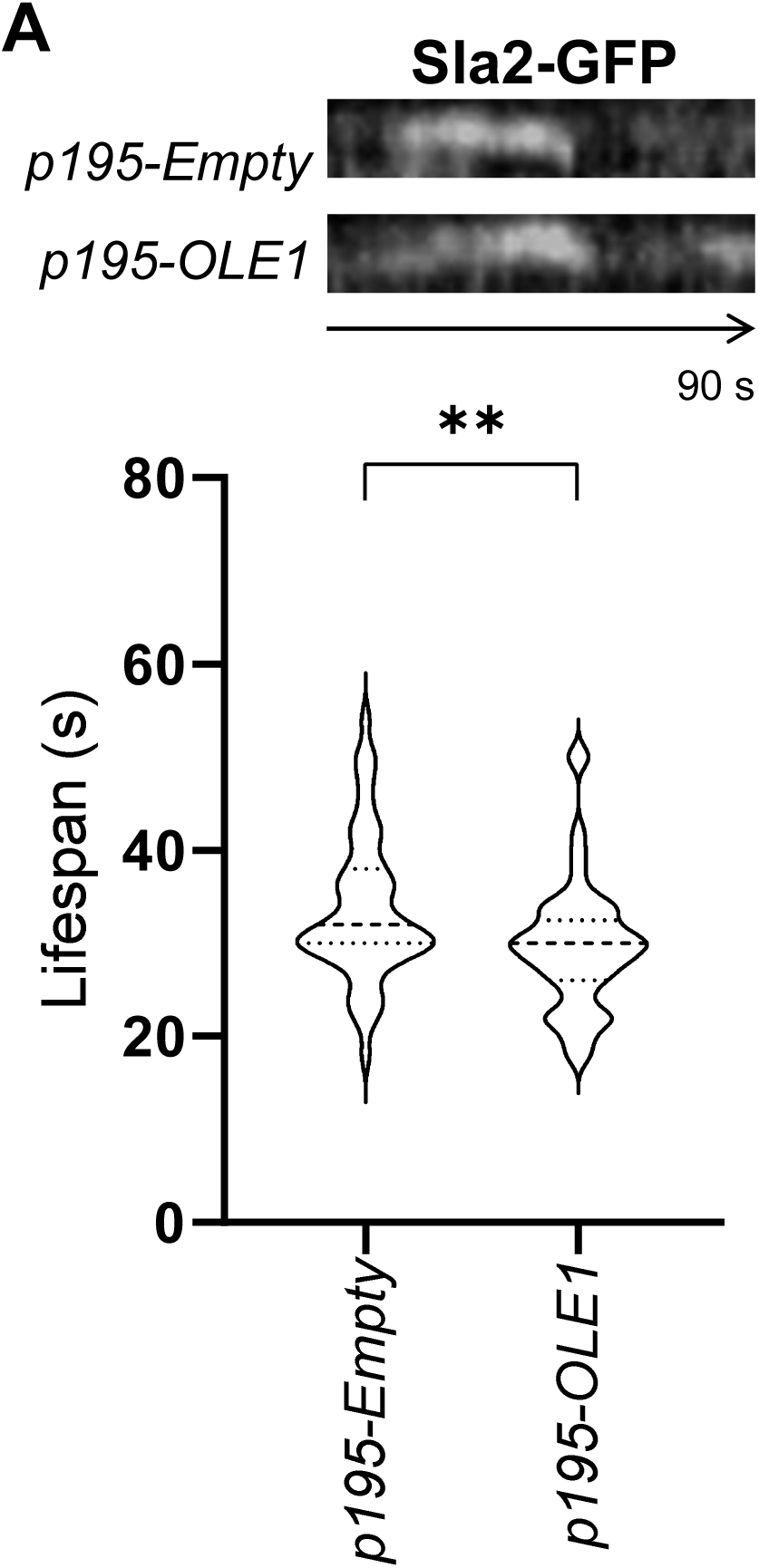
Overexpression of *OLE1* slightly Accelerates the Endocytic Coat Dynamics. Kymographs of 2 ss time-lapse epi-fluorescence movies of cortical Sla2-GFP patches for wt cells overexpressing *OLE1* (p195*OLE1*) from a multicopy plasmid or bearing the empty plasmid (p195). The violin plots show the Sla2-GFP cortical patch lifespans in the indicated strains. Asterisks indicate statistically significant differences between groups determined by Mann-Whitney test. Dashed lines indicate the median and dotted lines indicate the interquartile range (25^th^–75^th^ percentiles). n > 50 endocytic patches from at least 20 cells, from a representative experiment.

**Figure S4.**
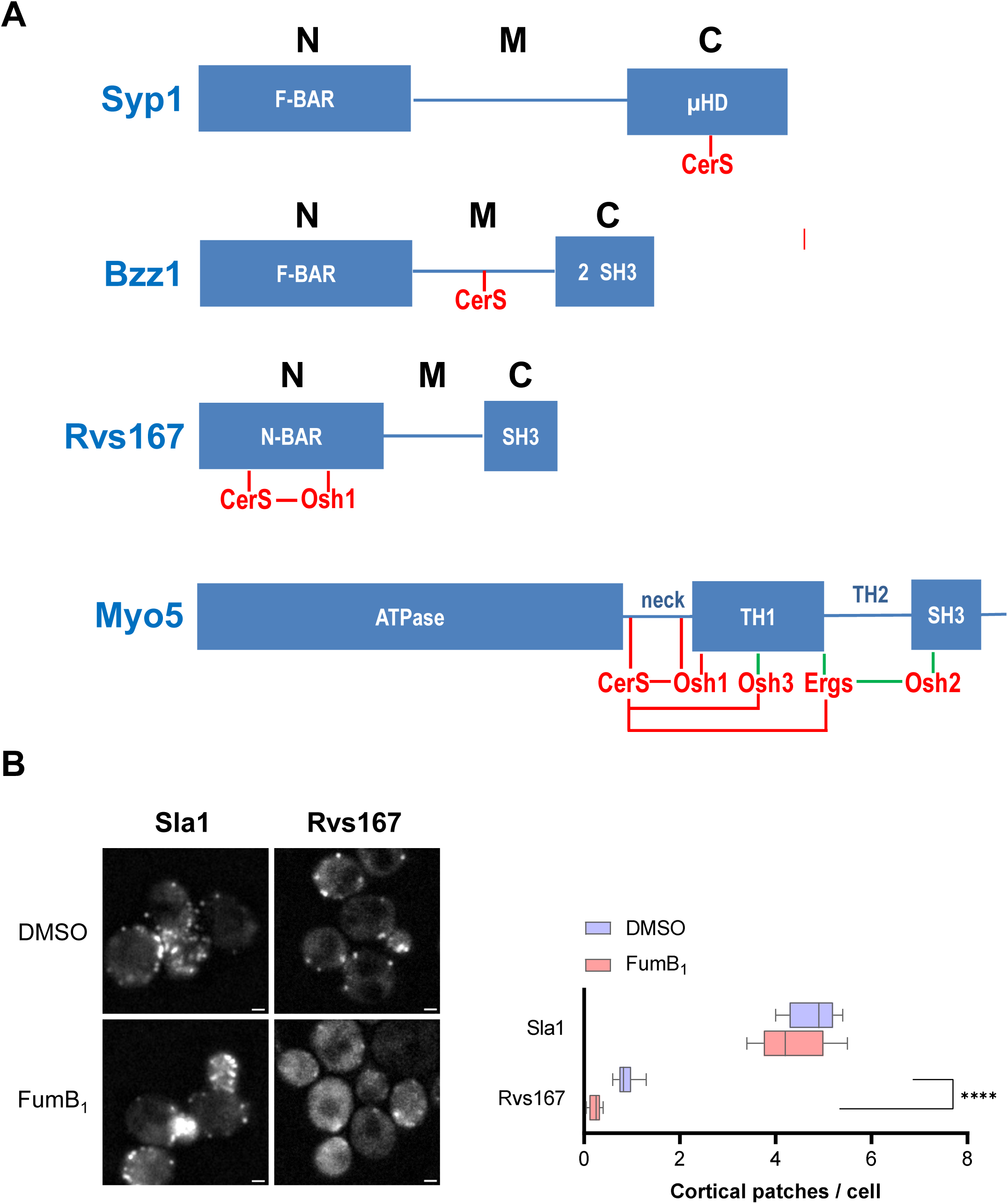
Rvs167-GFP Is not Recruited in Cortical Patches upon Inhibition of Cer Synthesis. **A.** Diagram of the interactions detected in this study (in red). In green, interactions demonstrated in (Encinar Del Dedo et al., 2021; Encinar Del Dedo et al., 2017). N, M and C indicate the N-terminal, middle, and C-terminal regions of the BAR (Bin/Amphiphysin/Rvs domains)-containing proteins; μHD, muniscin homology domain; SH3, Scr homology 3; TH1, tail homology domain 1; TH2, tail homology domain 2. **B.** Epi-fluorescence micrographs of live wt cells expressing either Sla1-GFP or Rvs167-GFP treated with 100 µM of Fumonisin B_1_ or mock treated; and box and whiskers plots of the number of the indicated cortical patches per cell in the corresponding experimental conditions. Boxes represent the 25^th^–75^th^ percentiles, the line indicates the median, and whiskers represent the 5^th^–95^th^ percentiles. n > 120 cells per strain, from a representative experiment. The number of endocytic patches was only quantified in the mother cells for clarity. Asterisks indicate statistically significant differences between groups determined by Welch’s t-tests. Bar = 1 μm.

**Figure S5.**
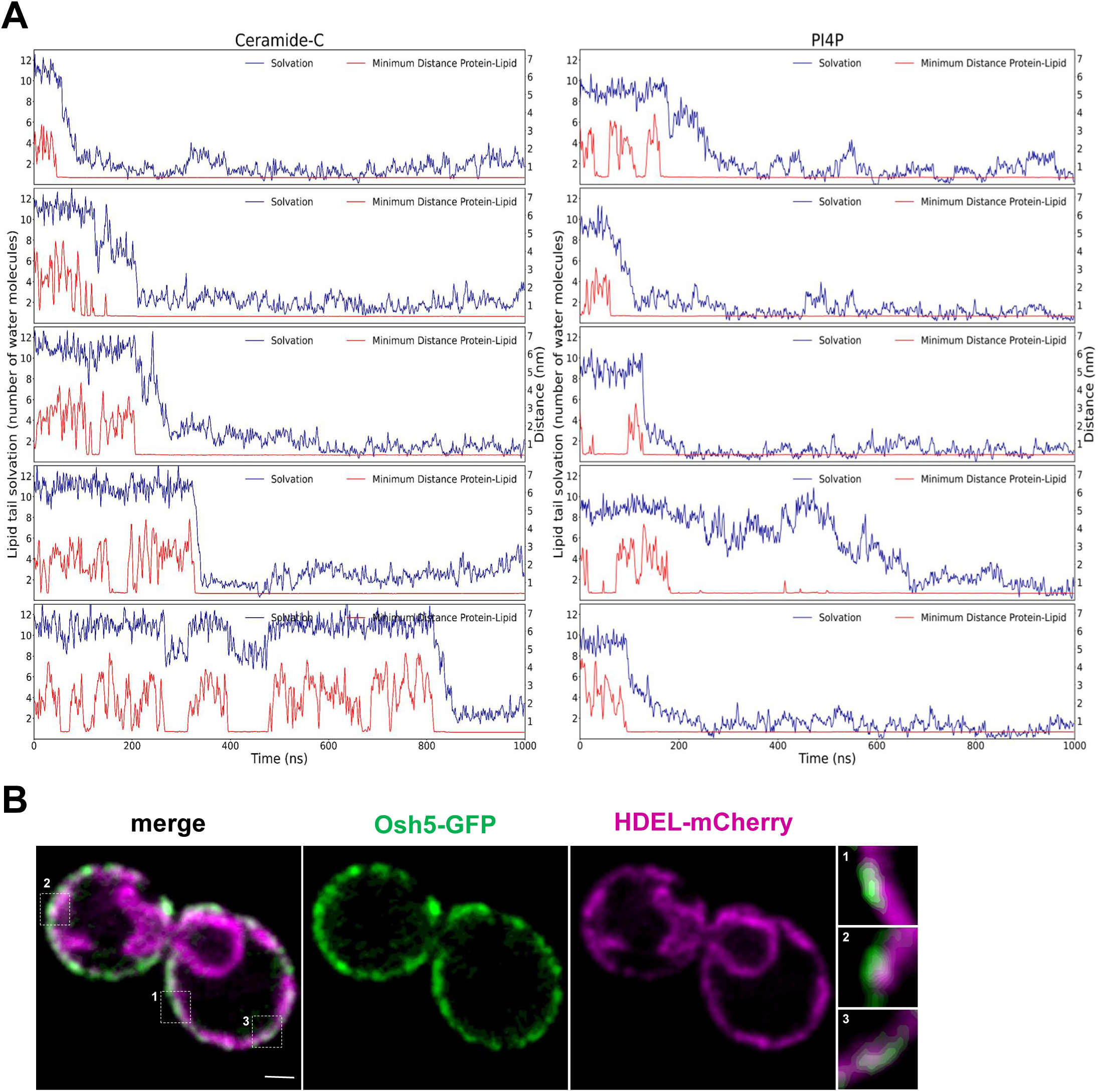
VLCFA-Cers Accommodate into the Hydrophobic Cavity of the Osh3 ORD. **A.** 1 µs time traces of minimum distance values between the lipids and Osh3 (red) along with the lipid tail solvation (blue). Both plots have been block-averaged every 10 frames. **B & C.** Dual color confocal-fluorescent micrographs of live yeast expressing Osh5-GFP and HDEL-mCherry. The individual channels of the same field and the merge are shown. Magnified micrographs of the depicted areas are shown on the right. Bar = 1 µm.

**Figure S6.**
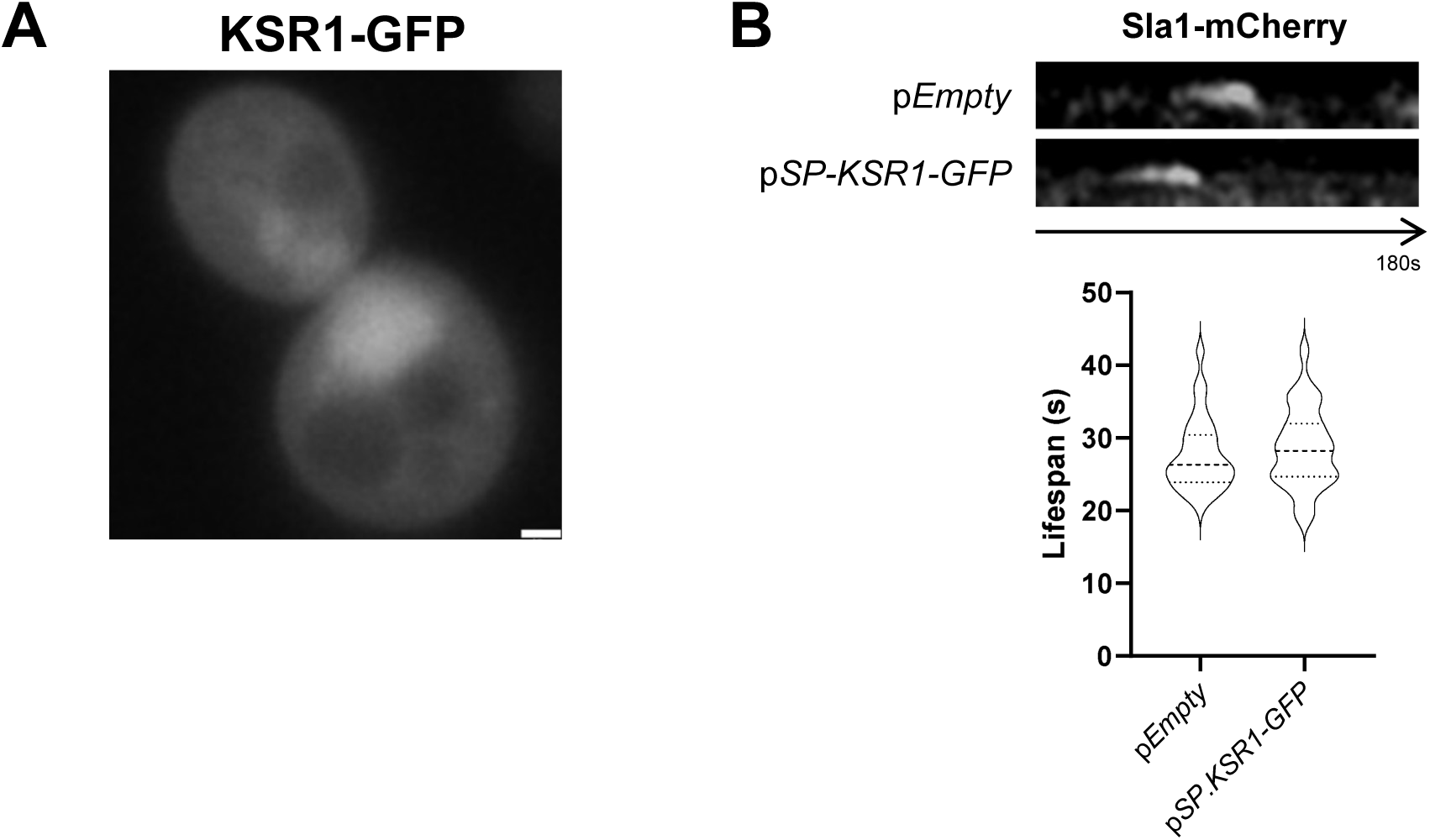
Endocytosis is not Altered by Expression of SP-KSR1-GFP. **A.** Epi-fluorescence micrographs of live yeast expressing KSR1-GFP. Bar = 1 µm. **B.** Kymographs of 2 s time-lapse epi-fluorescence movies of cortical Sla1-mCherry patches of wt cells expressing SP-KSR1GFP or bearing an empty plasmid. The violin plots show the lifespans of the Sla1-mCherry in the indicated strains. No statistically significant differences were detected by Mann-Whitney test. Dashed lines indicate the median and dotted lines indicate the interquartile range (25^th^–75^th^ percentiles). n > 50 endocytic patches from at least 20 cells, from a representative experiment.

## Notes

https://github.com/MolecularImagingPlatformIBMB/yeast-microscopy-pipeline

### Competing Interest Statement

The authors have declared no competing interest.

