## Supplementary figures for "Role of Very Long-Chain Fatty Acyl-Ceramides in Endocytic Uptake"

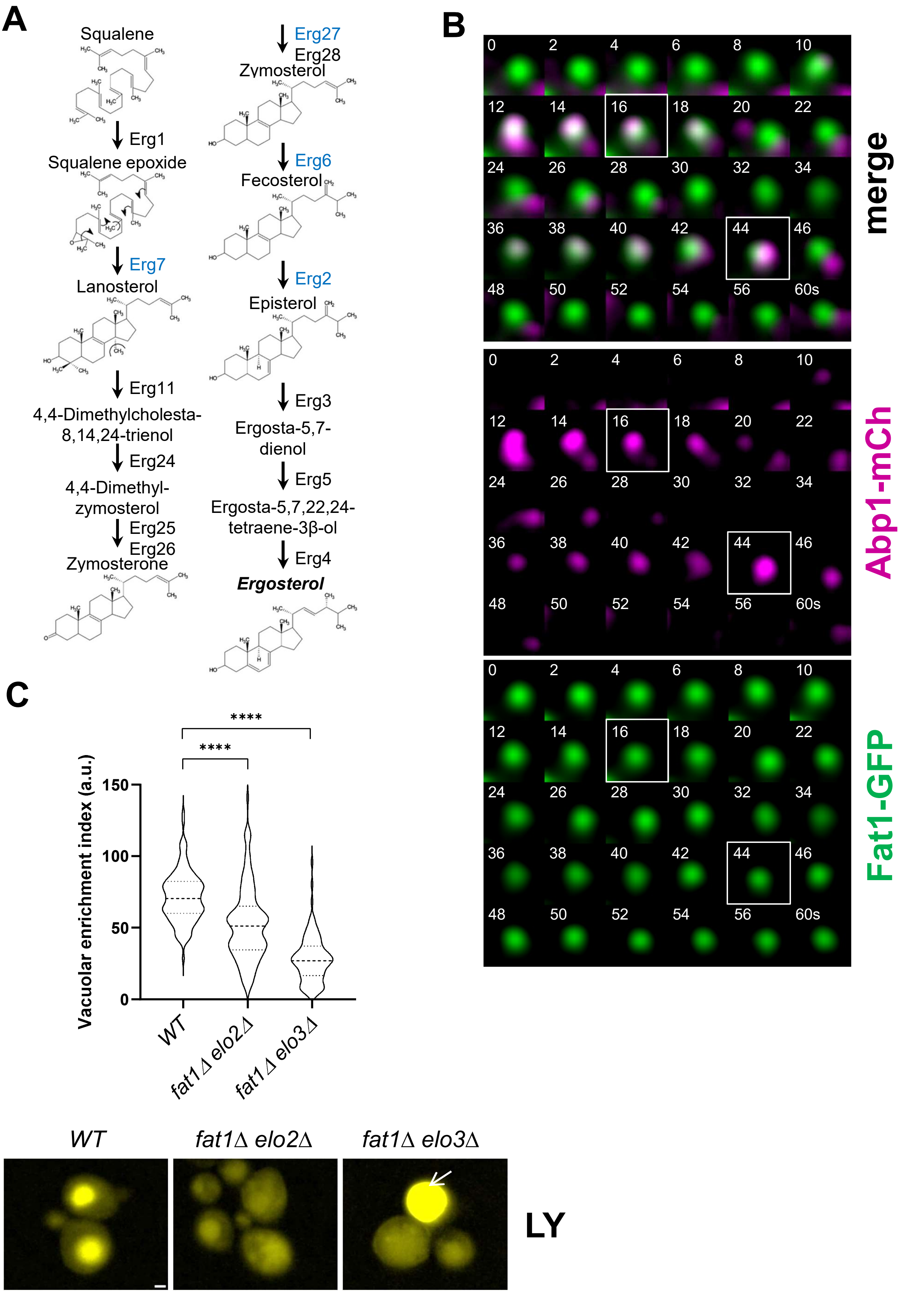
A
Squalene
Erg1
Squalene epoxide
Erg7
Lanosterol
Erg11
4,4-Dimethylcholesta-8,14,24-trienol
Erg24
4,4-Dimethyl-zymosterol
Erg25
Erg26
Zymosterone
Erg27
Erg28
Zymosterol
Erg6
Fecosterol
Erg2
Episterol
Erg3
Ergosta-5,7-dienol
Erg5
Ergosta-5,7,22,24-tetraene-3β-ol
Erg4
Ergosterol
B
merge
Abp1-mCh
Fat1-GFP
0 2 4 6 8 10
12 14 16 18 20 22
24 26 28 30 32 34
36 38 40 42 44 46
48 50 52 54 56 60s
C
Vacuolar enrichment index (a.u.)
0 50 100 150
****
****
WT
fat1Δ elo2Δ
fat1Δ elo3Δ
LY

Figure S1

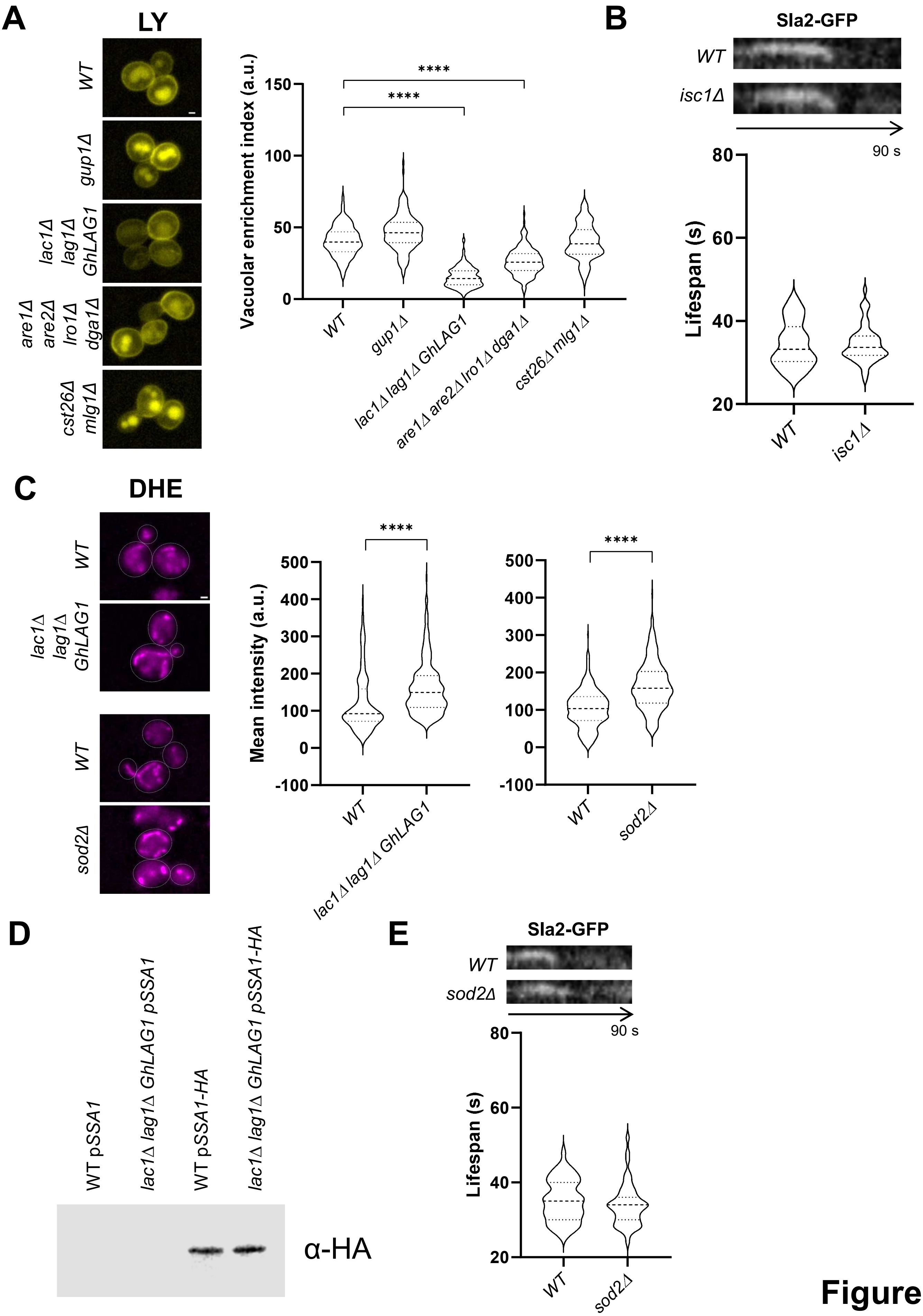
A
LY
WT
gup1Δ
lac1Δ lag1Δ GhLAG1
are1Δ are2Δ lro1Δ dga1Δ
cst26Δ mlg1Δ
Vacuolar enrichment index (a.u.)
****
****
150
100
50
0
WT
gup1Δ
lac1Δ lag1Δ GhLAG1
are1Δ are2Δ lro1Δ dga1Δ
cst26Δ mlg1Δ
B
Sla2-GFP
WT
isc1Δ
90 s
Lifespan (s)
80
60
40
20
WT
isc1Δ
C
DHE
WT
lac1Δ lag1Δ GhLAG1
WT
sod2Δ
Mean intensity (a.u.)
****
500
400
300
200
100
0
-100
WT
lac1Δ lag1Δ GhLAG1
****
500
400
300
200
100
0
-100
WT
sod2Δ
D
WT pSSA1
lac1Δ lag1Δ GhLAG1 pSSA1
WT pSSA1-HA
lac1Δ lag1Δ GhLAG1 pSSA1-HA
α-HA
E
Sla2-GFP
WT
sod2Δ
90 s
Lifespan (s)
80
60
40
20
WT
sod2Δ

**Figure S2**

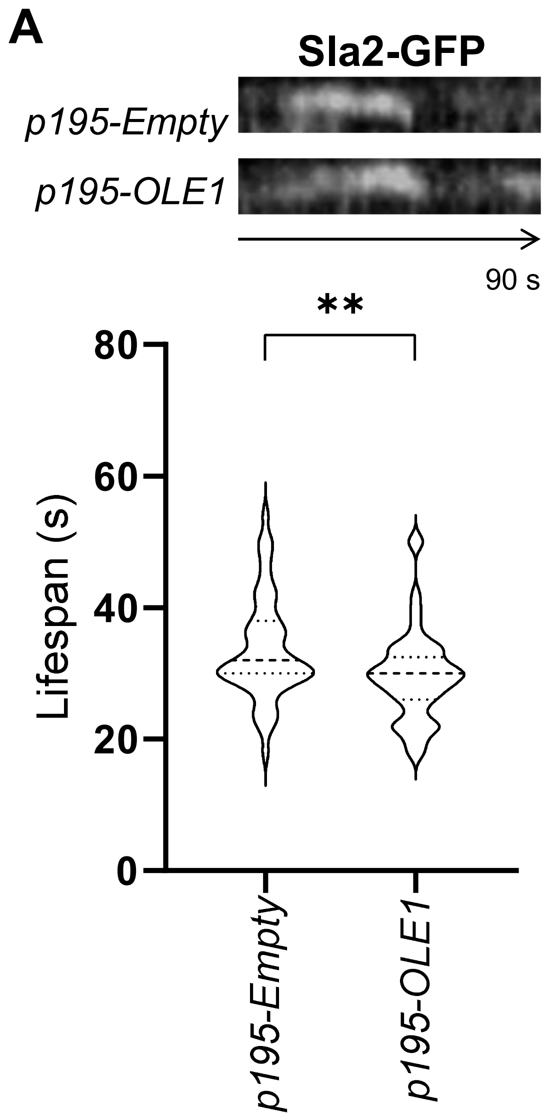
A
Sla2-GFP
p195-Empty
p195-OLE1
90 s
**
80
60
40
20
0
Lifespan (s)
p195-Empty
p195-OLE1

**Figure S3**

A

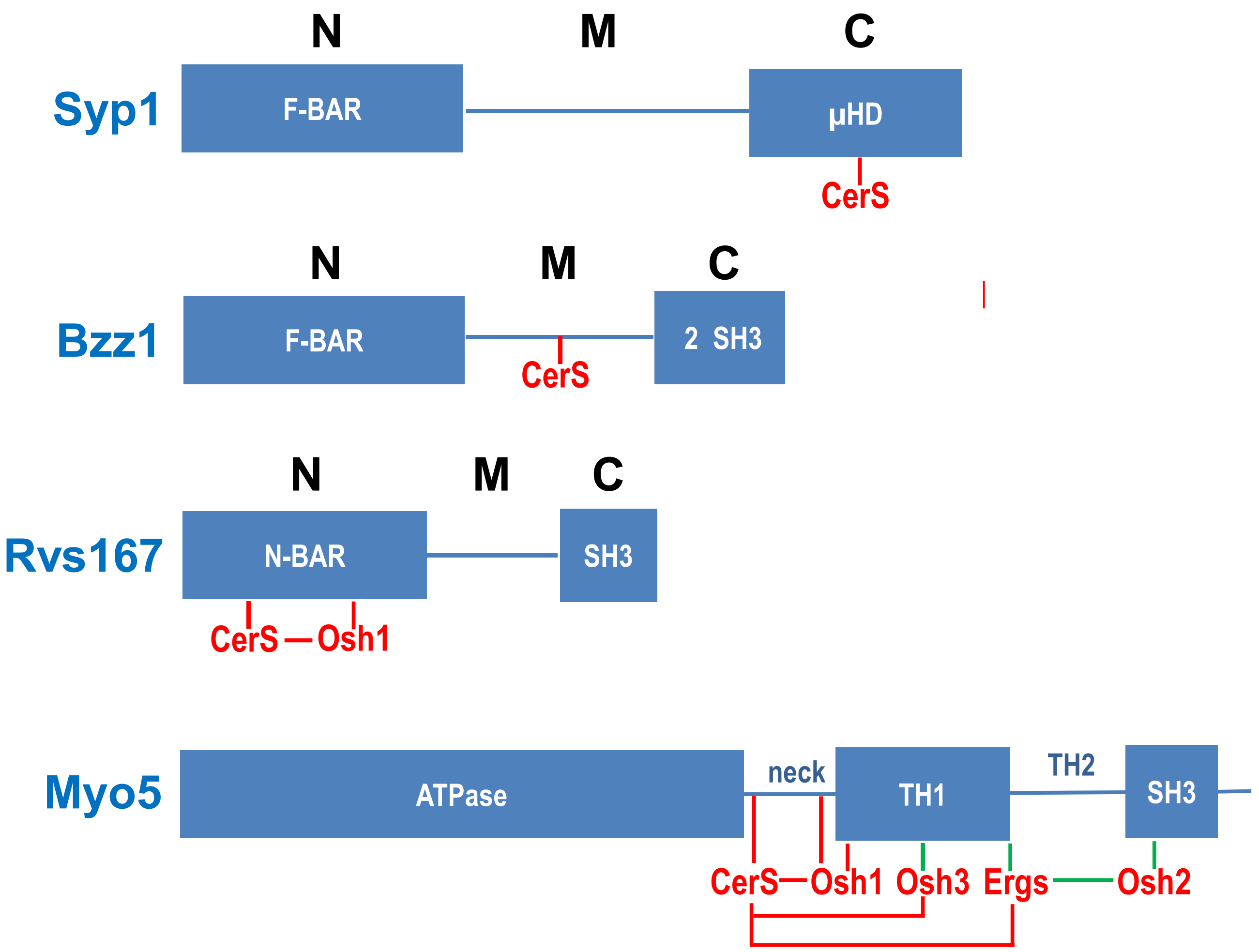
N
M
C
Syp1
F-BAR
μHD
CerS
N
M
C
Bzz1
F-BAR
2 SH3
CerS
N
M
C
Rvs167
N-BAR
SH3
CerS
Osh1
Myo5
ATPase
neck
TH1
TH2
SH3
CerS
Osh1
Osh3
Ergs
Osh2

B

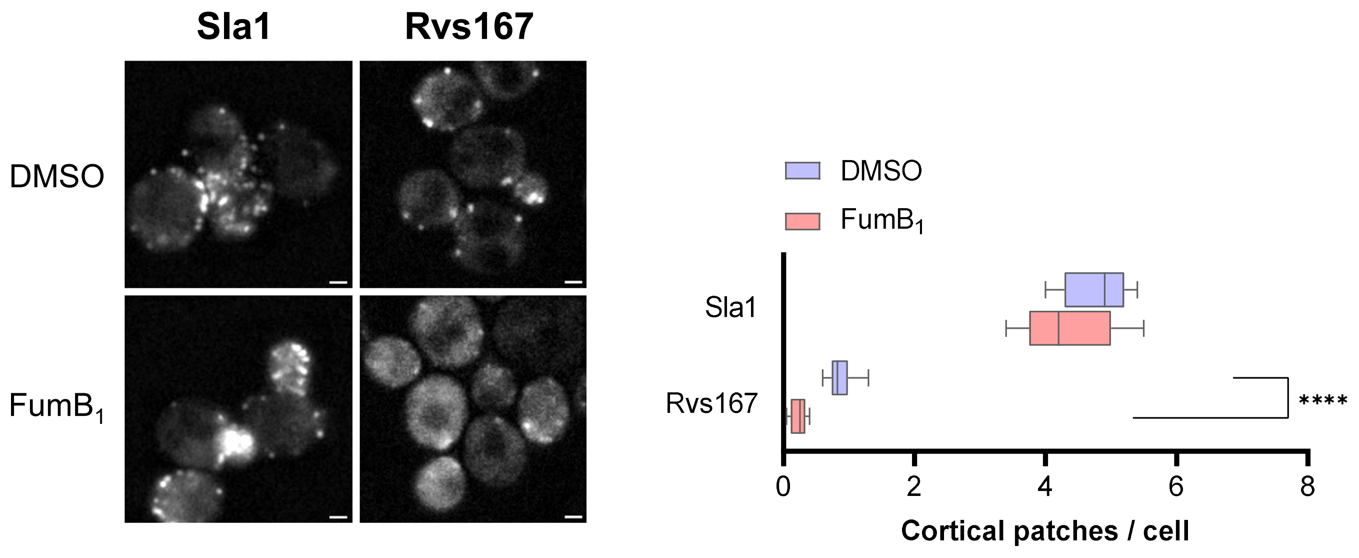
Sla1
Rvs167
DMSO
FumB1
DMSO
FumB1
Sla1
Rvs167
****
0
2
4
6
8
Cortical patches / cell

Figure S4

**A**

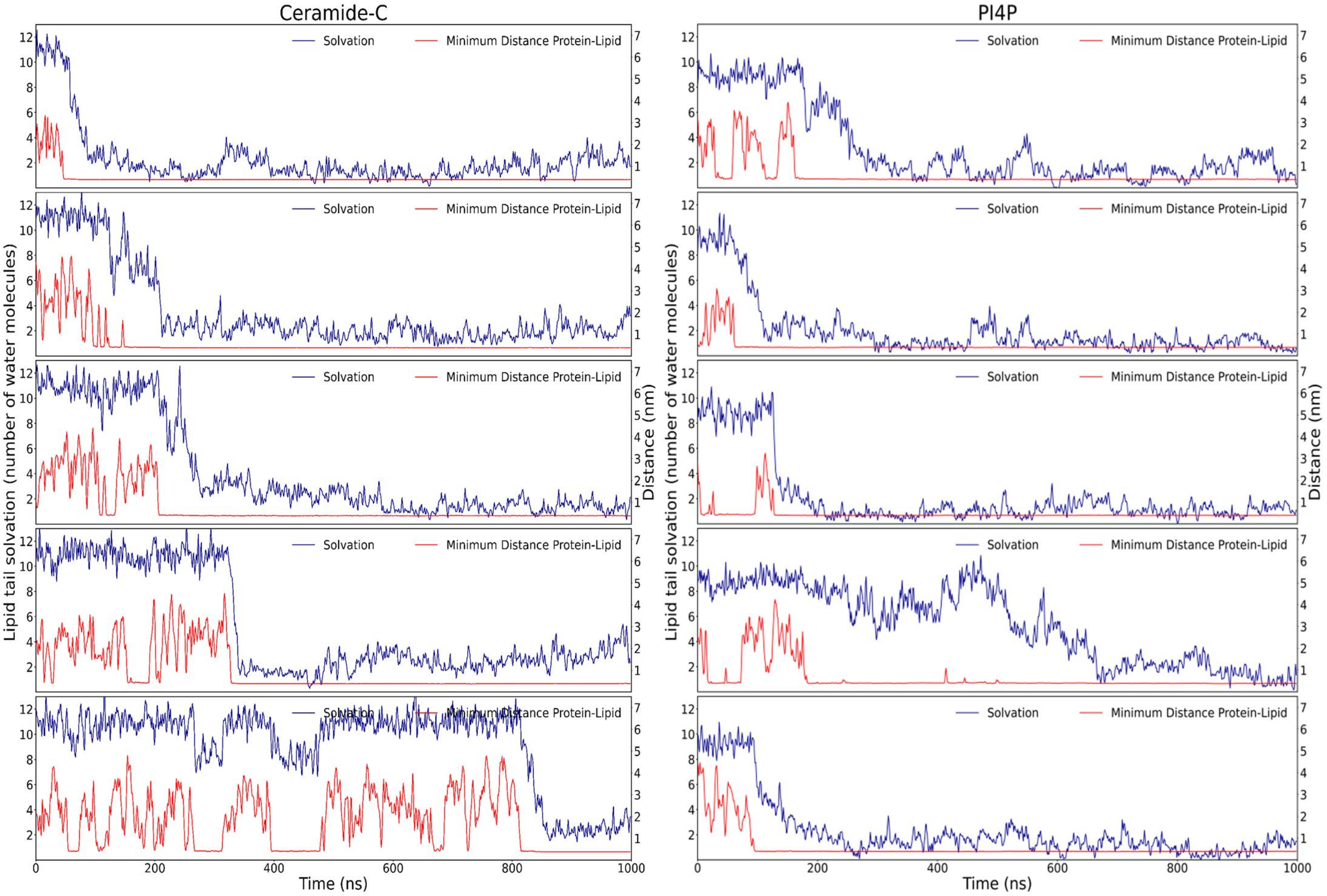
Ceramide-C
PI4P
Solvation
Minimum Distance Protein-Lipid
Lipid tail solvation (number of water molecules)
Distance (nm)
Time (ns)

**B**

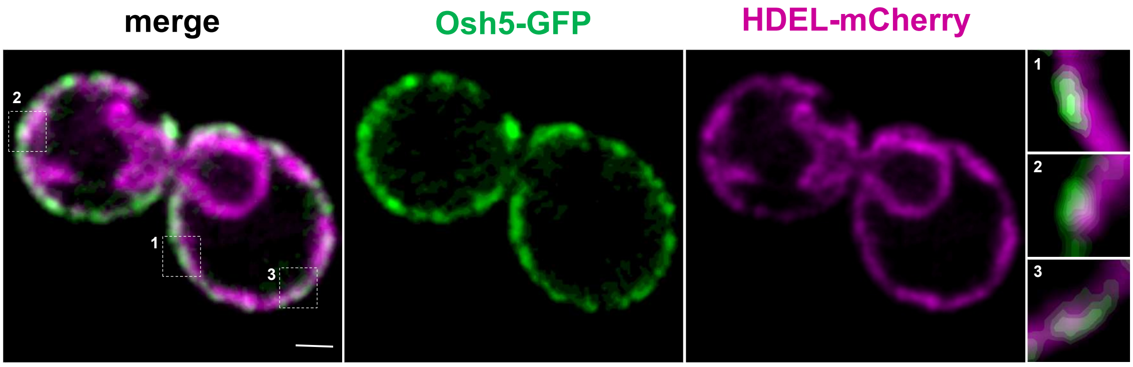
merge
Osh5-GFP
HDEL-mCherry
1
2
3

**Figure S5**

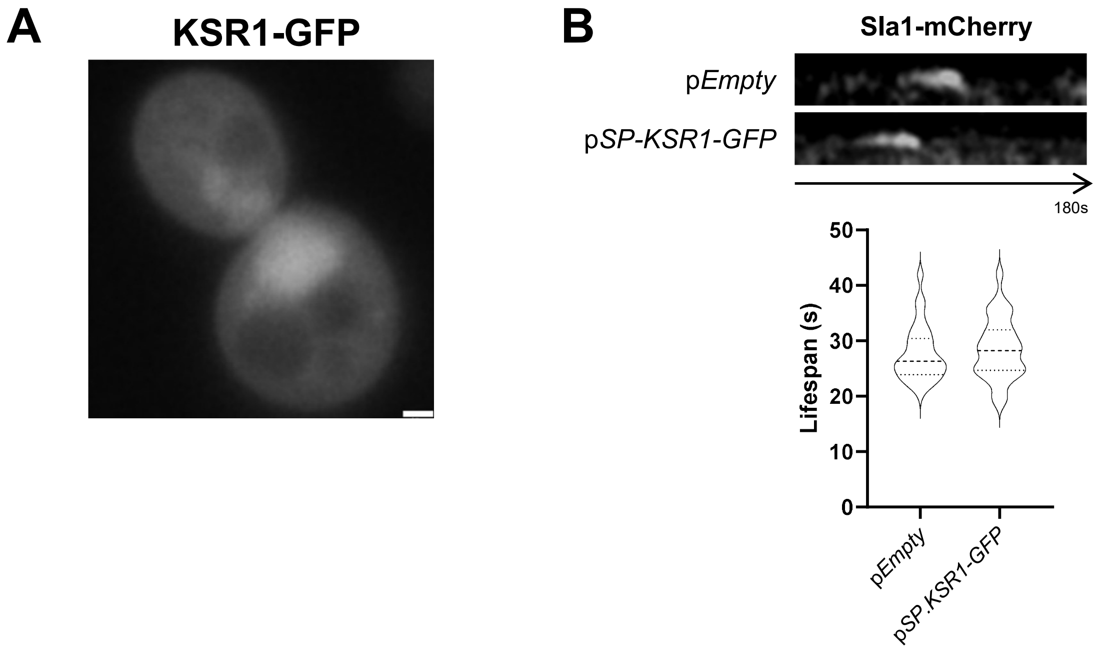
A
KSR1-GFP
B
Sla1-mCherry
pEmpty
pSP-KSR1-GFP
180s
Lifespan (s)
50
40
30
20
10
0
pEmpty
pSP.KSR1-GFP

**Figure S6**
